# Pedigree-based genome-wide imputation using a low-density amplicon panel for the highly polymorphic Pacific oyster *Crassostrea* (*Magallana*) *gigas*

**DOI:** 10.1101/2025.02.11.637717

**Authors:** Ben J. G. Sutherland, Konstantin Divilov, Neil F. Thompson, Thomas A. Delomas, Spencer L. Lunda, Christopher J. Langdon, Timothy J. Green

## Abstract

High-density genomic data are instrumental for selective breeding, but the costs associated with these approaches can hinder progress, as is the case for most aquaculture species. A strategy to reduce genotyping costs is to genotype a few select individuals at high-density (e.g., parents, grandparents), and many others at low density (e.g., offspring), then impute genotypes. This has been demonstrated *in silico* for Pacific oyster *Crassostrea* (*Magallana*) *gigas* but was particularly challenging relative to other species and has never been empirically tested. Here, four families of Pacific oysters, bred via marker-assisted selection for variation at a locus for field survival in an ostreid herpesvirus 1 (OsHV-1)-positive estuary, were exposed to OsHV-1 then genotyped using a low-density amplicon panel (n = 240 individuals). Parents were genotyped with the amplicon panel and by whole-genome resequencing. Offspring genotypes were imputed, and accuracy was determined by comparing against held-out whole-genome data for offspring. Imputation resulted in reduced minor allele frequencies and enriched homozygosity relative to empirical data. An *in silico* three-generation analysis was used to investigate the effect of deepening the pedigree, resulting in superior concordance in genotypes (GC = 84.5%) and allelic dosage (r = 0.73) compared to two-generation imputation (GC = 75.3%; r = 0.63). Genome-wide associations to OsHV-1 survivorship with imputed data identified significantly associated regions on the expected chromosome 8, but not at the expected position based on previous work, pointing to a potentially more complex genetic architecture for the trait. Our results empirically demonstrate the utility of amplicon panel-based genome-wide imputation in shellfish, and thus enable low-cost selective breeding techniques.

## Introduction

The Pacific oyster *Crassostrea* (*Magallana*) *gigas* (Thunberg, 1793) is a highly valuable aquaculture species globally (Botta et al. 2020). In many countries it is the subject of genetic improvement programs that range from large-scale governmental and commercial programs to small-scale university programs. Access to genomic resources for marker-assisted or genomic selection quickens economic gains and advances food production (Martínez-García et al. 2022) but is significantly limited by the availability of funding, which creates a need for cost-effective options (Boudry et al. 2021; Delomas et al. 2023). The recent development of a low-density Pacific oyster amplicon panel for SNP genotyping is anticipated to facilitate high-throughput lower-density genotyping using a repeatable and low-cost approach (Meek and Larson 2019; Sutherland et al. 2024). However, with a density of 592 amplicons across approximately 640 Mbp and 10 chromosomes (Peñaloza et al. 2021), alone it is limited in scope for methods requiring genome-wide coverage, especially given the rapid linkage decay in oysters (Hollenbeck and Johnston 2018; Hu et al. 2022; Mao et al. 2024).

Genome-wide coverage can be attained cost-effectively through imputation by combining low-density genotypes with high-density genotypes of select individuals (Habier et al. 2009), such as parents and grandparents (Delomas et al. 2023) and assuming Mendelian inheritance. This has been recently explored *in silico* within Pacific oysters. Kriaridou et al. (2023) subsampled SNPs from high density data for Pacific oysters and three vertebrate species to construct low density genotype samples, imputed genotypes, then determined imputation accuracy relative to empirical genotypes. This study demonstrated the challenge of obtaining high-quality imputation results that is specific to Pacific oysters when only including parents and offspring. Delomas et al. (2023) simulated genotypes from a breeding program and found that low-density panels (e.g., 250 or 500 SNP markers) combined with high-density genotypes from grandparents and parents allowed imputation to provide reliable genotypes for genomic selection (GS). However, empirical use of an amplicon panel and high-density data to conduct imputation has not yet been evaluated for any bivalve species, including the Pacific oyster.

Traits with unknown underlying causal mechanisms or polygenic architecture are not candidates for marker-assisted selection (MAS) but can be targeted by GS. Cost-effectiveness is necessary for GS in Pacific oysters and other aquaculture species (Delomas et al. 2023), especially if there are differences in phenotypic and genetic responses to local environmental conditions (Allen et al. 2020; Allen et al. 2021). Furthermore, the ability to target multiple traits is essential for Pacific oyster aquaculture, considering the challenges that the industry has experienced from biotic and abiotic stressors (Green et al. 2019; King et al. 2019). These stressors can result in summer mortality syndrome (Cowan et al. 2023; Cowan et al. 2024) and other massive mortality events, in some cases associated with changes in climate or extreme weather events (Raymond et al. 2022). Adverse environmental conditions may change over time, creating a moving target for breeding programs that requires affordability and adaptability. Resilience through selective breeding has been highlighted as an important adaptation strategy for both Pacific oysters and eastern oysters *C. virginica* (Langdon et al. 2003; de Melo et al. 2016; Melo et al. 2018; Allen et al. 2020; Green et al. 2023).

The field of shellfish genomics presents unique challenges, as has been observed in Pacific oysters. Segregation distortion and non-Mendelian transmission has long been observed in Pacific oysters (Launey and Hedgecock 2001), and has challenged genetic map construction, although this has been largely resolved (Yin et al. 2020) and chromosome-level genomes are now available (e.g., Peñaloza et al. 2021; Qi et al. 2021). Very high polymorphism levels in oysters (Hedgecock et al. 2005; Sauvage et al. 2007; Plough 2016) pose challenges for some genetic methods, for example due to the high frequency of null alleles (Hedgecock et al. 2004). Putative null alleles observed in a low-density amplicon panel impeded parentage assignments, necessitating their removal for optimal performance (Sutherland et al. 2024; Thompson et al. 2025). Similarly, Guo et al. (2023) used a two-step process to reduce the abundance of null alleles in a SNP chip for eastern oysters. Alternatively, high polymorphism levels can be beneficial to some genetic methods, for example by enabling identification of additional variants within amplicon sequences (e.g., microhaplotypes) to increase statistical power for relationship assignment and extending the utility of an amplicon panel to populations not included in the original panel design (Thompson et al. 2025). Additionally, these novel variants will increase power for imputation (Delomas et al. 2024).

In the present study, we used the Cgig_v.1.0 amplicon panel (Sutherland et al. 2024) for pedigree-based imputation using Pacific oyster families (parents and offspring) from the Molluscan Broodstock Program (MBP) of Oregon State University. These families were selectively bred for field survival in a bay with seasonal outbreaks of ostreid herpesvirus 1 (OsHV-1). Parents were genotyped at high density using a target of 20x coverage whole-genome resequencing. Offspring (n = 240) were genotyped with the 592 locus amplicon panel. The objectives of this study were two-fold: (1) to evaluate the accuracy of genome-wide imputation using the amplicon panel; and (2) to conduct a genome-wide association study (GWAS) using imputed offspring genotypes to identify regions of the genome associated with resistance to OsHV-1. The offspring samples (n = 240) were part of a laboratory OsHV-1 exposure trial and have directly observed survival phenotypes, but many of the mortalities had poor DNA quality for whole-genome resequencing, impeding the ability to conduct a GWAS for OsHV-1 survival. However, the DNA of these juveniles was of sufficient quality for amplicon panel genotyping, and therefore we used pedigree-based imputation with the amplicon panel genotypes to perform the GWAS. Offspring with sufficient high-quality DNA for whole-genome resequencing (n = 176) were not used for imputation but rather were used to evaluate imputation accuracy and to further evaluate differential characteristics of imputed genotypes. To evaluate the effect of deepening the pedigree on the accuracy of imputation, previously-published grandparent RAD-sequencing data (Divilov et al. 2023b) was used to create an *in silico* panel for all three generations to use for imputation, and compared with the two-generation results. Collectively these results provide insights on the reliability of pedigree and amplicon panel-based imputation for generating high-density imputed data for use in large- and small-scale Pacific oyster breeding programs and related analyses.

## Methods

### Pacific oyster families and OsHV-1 laboratory challenge

Offspring from four Pacific oyster families (F114-F117) from the Molluscan Broodstock Program (MBP) were used in an exposure trial to ostreid herpesvirus 1 (n = 60 offspring per family), as previously described (Lunda et al. *in review*). Cohorts and parent families are shown in Sutherland et al. (2024). The eight parents of the four families were heterozygous for the OsHV-1 survival/ resistance marker on chromosome 8 (Divilov et al. 2023b; Lunda et al. *in review;* Surry et al. 2024). Each exposed animal had phenotypes recorded including the survival state by the end of the 16-day trial (dead or alive) and the day post-exposure (DPE) on which the mortality occurred. To differentiate individuals that died on the final day of the trial (day 16), individuals that survived until the end of the trial (i.e., ‘alive’) were assigned a value of 17 DPE for the mortality date. All samples were preserved in DNA/RNA Shield (Zymo Research) upon detection of mortality (open valves) or moribundity (no response to tactile stimulation) during daily tank inspections.

### DNA extraction and library prep

Genomic DNA was extracted from offspring (n = 240) and parents (n = 8), and quantity and quality were assessed by fluorescence (Qubit) and 1% agarose gel electrophoresis, respectively. Offspring that passed quality evaluations were used for whole-genome resequencing in paired-end 150 bp reads to a target depth of 10x, and all parents were sequenced to a target depth of 20x on a NovaSeq sequencer (Illumina). Extractions and library preparations were conducted by the Centre for Aquaculture Technologies (CAT), San Diego, California (USA).

### High-density genotyping of parents and offspring

Whole-genome resequencing reads were analyzed using the pipeline *wgrs_workflow* (see Data Availability). Samples were checked for quality using FastQC and MultiQC (Andrews 2010; Ewels et al. 2016), trimmed with cutadapt (v.3.5; Martin 2011) in paired-end mode to remove Illumina adapters (maximum adapter error rate = 20%), low-quality bases, and reads smaller than 50 bp (flags: -q 15 –trim-n and -m 50), and then re-inspected for quality. Trimmed reads were then aligned to a chromosome-level Pacific oyster reference genome (GCF_902806645.1; Peñaloza et al. 2021) using bwa mem (Li 2013) with read group identifiers added per sample. SAMtools (Li 2011) was used to convert SAM files to BAM format, filter for quality (-q 10), as well as to sort and index the alignment files. Approximate sequencing depth was calculated based on a genome size of 640 Mbp (Peñaloza et al. 2021). Duplicate alignments were marked and tallied using the Picard Toolkit (Broad Institute 2024). Reads from all samples (offspring and parents) were merged into a single BAM file using SAMtools in preparation for genotyping.

Genotyping was conducted with bcftools mpileup (Danecek et al. 2021) with base alignment quality (BAQ) applied (-D) and a maximum depth (-d) per merged BAM file of 11,500 reads. Variants were annotated by mpileup with allelic depth and total depth, and bcftools call was used in multiallelic caller mode (-m) to output variant sites (-v) and annotate genotype quality (--annotate GQ) to a BCF file. The BCF file was filtered using bcftools to remove variants that were within 5 bp of an indel and those with more than 10% missing data across all individuals. Only biallelic SNP variants were retained, and these required an overall quality score greater than 99, and an average depth across all samples of at least 10 reads. For each individual, any genotyped sites with fewer than five or more than 100 reads, or with per-site genotype quality of less than 20 were set to missing, and then the missing-data filter was re-applied across individuals, as above. The filtered data were then screened to remove any locus with a minor allele frequency (MAF) < 0.05, considering all genotyped parents and offspring together. This MAF-filtered dataset was used for all downstream applications. Whole genome genotypes comprised the high-density (HD) dataset. The HD dataset for parents was used for downstream imputation, and the HD dataset for offspring was held out from imputation and only used to evaluate imputation accuracy (Figure 1; see below). The HD datasets were also used to check genotype quality based on parent-offspring trios (see below), but this was not used during imputation.

**Figure 1.**
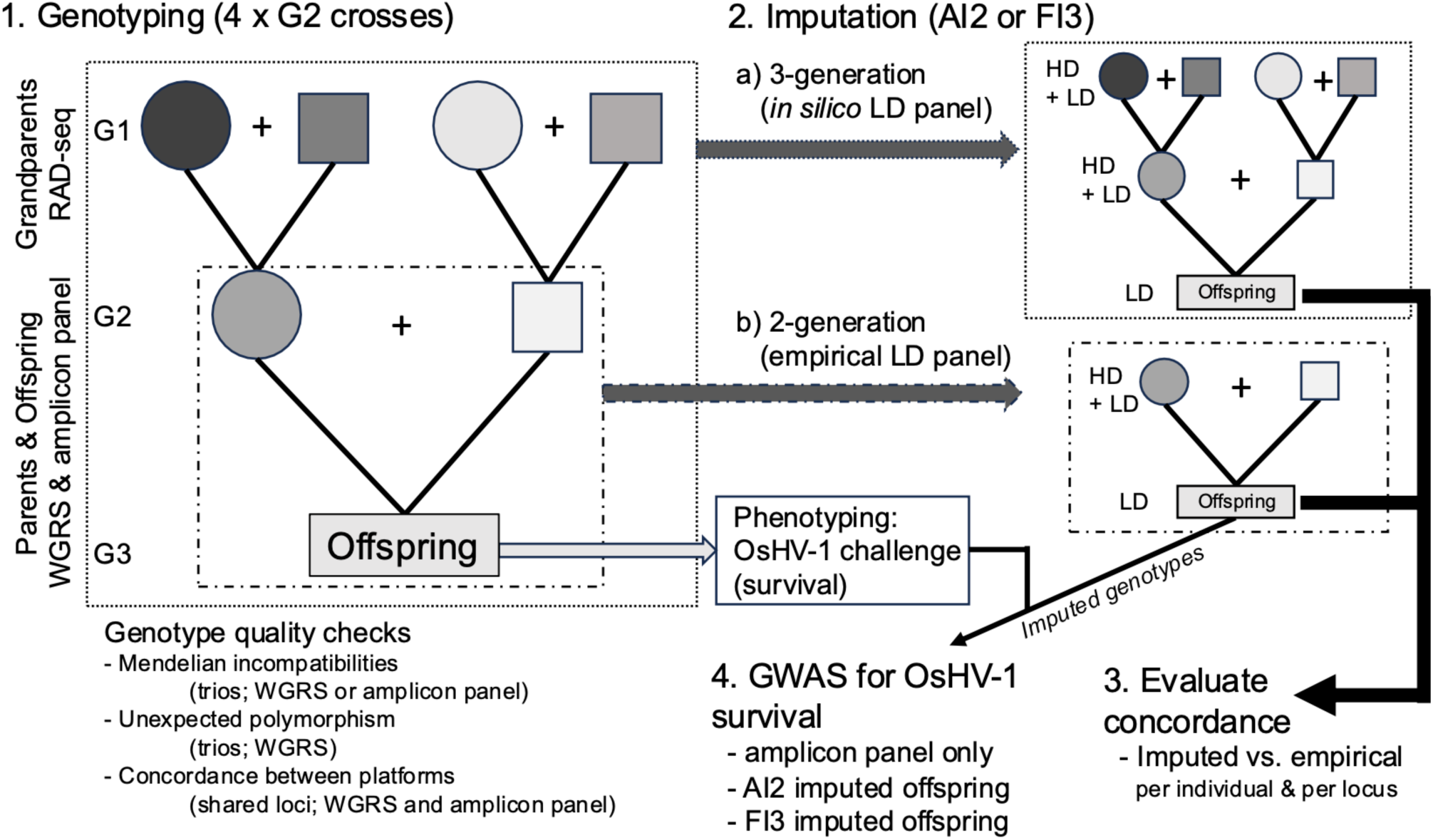
Workflow overview considering three- or two-generation (G1-G3) datasets and imputation by AlphaImpute2 (AI2) or FImpute3 (FI3). Dams are shown as circles and sires as squares. The imputation outputs from all datasets were evaluated for accuracy by comparisons with empirically genotyped offspring, but only the two-generation imputed genotypes were used as an input for a genome-wide association study for OsHV-1 survival.

### Amplicon panel library preparation, sequencing and genotyping

Genomic DNA samples from the 240 offspring were normalized and used for AmpliSeq library preparation (Thermo Fisher) with the *Cgig_v.1.0* amplicon panel (Sutherland et al. 2024) (SKU A58237 AGRISEQ PACIFIC OYSTER PANEL) by the Center for Aquaculture Technologies (San Diego). Parents (n = 8) were previously sequenced using the same panel (Sutherland et al. 2024). Output fastq or bam files were provided for the offspring and parents, respectively, and bam files were converted to fastq using the bamtofastq function of bedtools (Quinlan and Hall 2010). If replicate samples were present, the replicate with the highest sequencing depth was retained. Raw fastq data were aligned to the chromosome-level reference genome (GCF_902806645.1) using bwa mem, assigning read group identifiers and quality filtering to maintain alignments with MAPQ > 1. TorrentSuite (Thermo Fisher) VariantCaller VCF files were also obtained with the *Cgig_v.1.0* hotspot and regions file.

Aligned samples were used to genotype target and novel variants *de novo* (see Data Availability, *amplitools*). Alignment files were merged using SAMtools, and genotyping was conducted using bcftools mpileup and call (as above but with mpileup -d set to 115,000 reads), annotating the BCF file with allelic information and genotype quality values. Filtering of variants was conducted as described above for whole-genome resequencing data, except that the missing data filter allowed up to 15% missing data across samples, and the minimum and maximum depth cutoffs per genotype of 10 reads and 10,000 reads, respectively. The MAF-filtered dataset was used for all downstream applications. Amplicon panel genotypes were used in a principal component analysis (PCA) using the function *glPCA* of adegenet (v.2.1.5; Jombart and Ahmed 2011) in R (R Core Team 2025) to confirm clustering of families. The amplicon panel *de novo* genotypes for parents and offspring are referred to as the LD panel and were used for downstream imputation (Figure 1; see below).

### Identification of Mendelian incompatibilities, unexpected polymorphism, and platform concordance

Using the LD panel or HD datasets, parent and offspring trios were used to identify loci displaying Mendelian incompatibilities (MI) in each platform to evaluate genotype quality. MI were identified using the Mendelian function of bcftools with a pedigree file of expected trios, and MI were tallied per individual and per locus. Parent-offspring trios were also used in the HD dataset as an additional genotype quality check to identify loci with unexpected polymorphism in offspring when parents were monomorphic.

As an additional genotype quality check, the concordance of genotypes for loci present in both the LD panel and HD dataset was evaluated. Shared loci were identified using bcftools isec, and compared using bcftools stats, including the *verbose* flag for per-site concordance evaluation. Custom R scripts within the *impute_workflow* repository (see Data Availability) were used to inspect the results of bcftools stats, including the proportion of concordant genotypes per individual and per locus and the Pearson correlation of allelic dosage between the two platforms per individual.

### Grandparent genotype data and in silico panel construction

Grandparent RADseq sequence data for the families used in this study (F114-F117) were obtained from NCBI SRA (Divilov et al. 2023b) (see Data Availability), aligned to the reference genome (GCF_902806645.1), and genotyped as described above with bcftools mpileup and call. Family identifiers for each parent of families F114-F117 were provided by Sutherland et al. (2024; see Table 4). However, grandparents were not genotyped using the LD amplicon panel (genomic DNA from these individuals was not available at the time of sample preparation). Therefore, an *in silico* LD panel was constructed from available sequence data (Figure 1) to evaluate the effect of having three generations in imputation. To construct the *in silico* LD panel, shared loci between grandparents (RAD-seq data), parents (whole-genome resequence data), and offspring (whole-genome resequence data, n = 176 offspring) were identified using bcftools isec (with flag –collapse none). All shared loci were retained for the grandparents and parents, and an LD panel was created for the offspring by subsetting the shared loci to only retain a random selection of 1,000 loci (all other loci were set to missing; Figure 1).

The three generation *in silico* dataset was used for imputation to determine the effect of three generations on imputation accuracy (see below). For comparison with the empirical, panel-based two-generation dataset, the *in silico* dataset was also subset to only include the parents (HD data) and offspring (LD *in silico* panel) and imputed to estimate the accuracy of imputation with two generations. In parallel, the regular empirical dataset with the full HD parents and LD amplicon panel offspring was subsampled to only include 1,000 LD panel loci and a similar number of HD loci as the *in silico* dataset above to emulate the *in silico* dataset in the ratio of HD and LD loci for a full comparison. Accuracy was only evaluated on imputed loci by excluding LD panel loci prior to accuracy evaluation using bcftools isec (see below for Accuracy evaluation).

### Imputation

To construct the empirical imputation input dataset, the LD amplicon panel data for parents and offspring were combined with the HD dataset (whole-genome resequencing) from the parents. To combine the LD and HD datasets, all LD loci that were also present in the parent HD data were removed from the HD data using the bcftools isec function (with flag –*collapse all* to remove loci regardless of allelic identity). The LD and HD parent data were then combined using bcftools concat. The parent HD+LD data were then combined with the LD offspring data using bcftools merge (Figure 1).

The BCF file containing imputation input data (parent HD+LD; offspring LD) was converted to AlphaImpute2 format (Whalen and Hickey 2020) using custom R scripts, and a pedigree file was prepared. The ten Pacific oyster chromosomes were separated into individual files, AlphaImpute2 was run on each chromosome individually, and the results were combined into a single file. The data were similarly prepared for FImpute3 (Sargolzaei et al. 2014) to be run on individual chromosomes. The analysis pipeline with all custom scripts is provided (see Data Availability, *impute_workflow*). The FImpute3 imputation process was also conducted for the three- and two-generations *in silico* panel, and the equal ratio HD and LD loci empirical dataset (see above).

### Imputation and empirical data concordance

Imputed genotypes per chromosome were combined into a single file and converted to a VCF file using custom scripts (see Data Availability). The HD data for the offspring (not used in imputation) were limited to only include loci in common with the imputed data and then directly compared using bcftools stats and summarized and visualized using custom scripts. Loci present in both imputed and empirical data were also used to compare per-locus allele frequencies and calculate per-sample genotype proportions (i.e., proportion of homozygous reference, heterozygous, homozygous alternate, and missing; Figure 1).

### Genome-wide association study (GWAS)

Three offspring genotype datasets were used to conduct GWAS analyses based on survival to the OsHV-1 laboratory challenge: (1) amplicon panel, *de novo* loci (LD data) only; (2) LD data imputed to high density by AlphaImpute2; and (3) LD data imputed to high density by FImpute3. Imputed data were all based on the two-generations (i.e., parents and offspring), as this was the dataset with the most offspring samples with LD genotype information (from the amplicon panel). Amplicon panel or imputed genotypes were converted to GEMMA input formats (Zhou and Stephens 2012) using custom scripts (see Data Availability, *ms_cgig_chr8_oshv1* and *impute_workflow*, respectively). GEMMA was used to calculate kinship between samples and then to calculate associations per locus to the binary survival phenotype. Analyses were restricted to loci with MAF > 0.05. Manhattan plots were generated using fastman (Paria et al. 2022), highlighting LD loci, or using custom scripts. Feature annotations were obtained from the NCBI FTP site for Annotation Release 102 for the GCF_902806645.1 genome, and annotations within 10 kbp of regions of interest for the *de novo* panel only, or AlphaImpute2- or FImpute3-based GWAS results were collected into a table.

## Results

### High-density (HD) genotyping of parents and offspring by whole-genome resequencing

Parents (n = 8) from the Molluscan Broodstock Program “chromosome 8” selected families F114-F117 were sequenced to a target depth of 20x coverage. Parents had a per-sample average (± s.d.) number of read pairs of 67.4 ± 44.9 M (range = 32.7-174.7 M; Additional File S1). This equates to an approximate coverage of 31.6 ± 21.0x (range = 15.3-81.9x). All samples had at least 98.9% reads passing quality trimming. Aligning against the reference genome resulted in an average (± s.d.) per-sample alignment rate of 77.4 ± 0.7%. Parents had an average (± s.d.) duplicate read percentage of 19.0 ± 1.4% (range: 17.0-21.2%) based on estimates from Picard Toolkit (Additional File S1).

Offspring of the four families were to be sequenced to a target depth of 10x coverage by whole-genome resequencing. However, a group of 64 offspring samples (26.7% of all 240 offspring) did not have sufficient high-quality DNA for library preparation due to DNA degradation, and the degradation was often associated with mortality during the OsHV-1 challenge. For the samples that were sequenced (n = 176), offspring had an average (± s.d.) of 23.0 ± 5.2 M read pairs (range = 3.2-33.9 M; Additional File S1), which is approximately 10.8 ± 2.4x depth per individual (range = 1.5-15.8x; n = 24 samples with less than 10x coverage). Sample alignment rates (± s.d.) were on average 74.9 ± 7.7% and the estimated average percent duplication of read pairs was 15.3 ± 1.6% (range: 11.8-18.9%; see Additional File S1).

Variant calling was conducted using all parents and offspring together (n = 184 samples), and this resulted in the identification of 58,941,933 raw variants. Filtering was conducted in a stepwise process (Table S1), resulting in 424,951 biallelic SNPs (MAF > 0.05) being retained after all filters were applied. Per offspring averages of the genotypic states homozygous reference, heterozygous, and homozygous alternate were 262.6k (61.8%), 114.7k (27.0%), and 13.8k (3.2%), respectively (Table 1; Additional File S2). Parents had similar values (Table 1). Percentages of missing data for offspring and parents were on average 8.0% (33.9k) and 8.2% (34.9k), respectively. One parent, 58-9F (dam of F117) had a very high percentage of missing data (45.0% missing; Additional File S3 and Additional File S2).

**Table 1.** Average (± s.d.) per-sample genotype counts (k = thousand loci) and percentages of total loci for (A) offspring or parents in the empirical data (whole-genome resequencing, HD data; n = 176 samples); and (B) in the offspring imputed data by AlphaImpute2 (AI2) or FImpute3 (FI3; n = 240).

|  | **Offspring,**<br>**empirical** |  | **Parents,**<br>**empirical** |  |
| --- | --- | --- | --- | --- |
|  | **Counts (k)** | **(%)** | **Counts (k)** | **(%)** |
| *REF/REF* | 262.6 ± 49.5 | 61.8 ± 11.6 | 262.3 ± 38.3 | 61.7 ± 9.0 |
| *REF/ALT* | 114.7 ± 22.9 | 27.0 ± 5.4 | 113.3 ± 27.1 | 26.7 ± 6.4 |
| *ALT/ALT* | 13.8 ± 2.9 | 3.2 ± 0.7 | 14.6 ± 1.8 | 3.4 ± 0.4 |
| *Missing* | 33.9 ± 73.7 | 8.0 ± 17.3 | 34.9 ± 63.4 | 8.2 ± 14.9 |
| total | 425.0 | - | 425.0 | - |

(B)
|  | **Offspring,**<br>**imputed (AI2)** |  | **Offspring,**<br>**imputed (FI3)** |  |
| --- | --- | --- | --- | --- |
|  | **Counts (k)** | **(%)** | **Counts (k)** | **%** |
| *REF/REF* | 341.6 ± 3.3 | 85.0 ± 0.8 | 345.0 ± 3.9 | 85.8 ± 1.0 |
| *REF/ALT* | 27.8 ± 5.6 | 6.9 ± 1.4 | 28.2 ± 4.1 | 7.0 ± 1.0 |
| *ALT/ALT* | 32.8 ± 2.3 | 8.2 ± 0.6 | 28.9 ± 0.4 | 7.2 ± 0.1 |
| *Missing* | 0 | 0 | 0 | 0 |
| total | 402.1 | - | 402.1 | - |

To check genotype quality, polymorphic loci in the offspring were inspected for polymorphism in their respective parents. An average of 360,877 loci per family had no missing data in the parents; of these loci, parents and offspring had an average of 46.7% and 47.8% polymorphic loci, respectively (Table S2). An average of 2.2% of the loci that were polymorphic in the offspring were unexpected due to monomorphism in their respective parents (Table S2). Per family, loci that were polymorphic in both the offspring and parents followed a distribution with peaks at 0.25, 0.5, and 0.75 alternate allele frequencies (Figure S1A), whereas loci that were polymorphic in offspring and monomorphic in parents were largely low allele frequency variants (Figure S1B).

### Low-density (LD) genotyping of parents and offspring with the amplicon panel

The parents (n = 8) and offspring (n = 240) genotyped by the amplicon panel had on average (± s.d.) 0.81 ± 0.42 M and 0.14 ± 0.14 M reads per sample, respectively. One parent, 65_8F (the dam of F115), had significantly fewer reads than the other samples, with only 0.03 M reads obtained. Alignment rates per sample against the reference genome (GCF_902806645.1) for parents and offspring data were on average (± s.d.) 94.4 ± 0.5% and 91.8 ± 6.3%, respectively. The per-sample read and alignment counts are available in Additional File S1. Genotyping *de novo* resulted in an initial 218,925 raw variants before any filters; after stepwise filtering, 1,257 SNPs were identified (MAF > 0.05; Table S1). The removal of individuals with a low genotyping rate (GR; GR ≤ 70%) resulted in the retention of 214 out of the 234 individuals with phenotypic data (Figure S2A). After the low-GR samples were removed, loci had on average 2.0% missing data (Figure S3A), and all loci were polymorphic. A PCA using multi-locus genotypes showed individual clustering by family (Figure S4). This is hereafter referred to as the LD dataset.

The standard genotyping pipeline for a ThermoFisher amplicon panel (i.e., with the Torrent VariantCaller; TVC) identified 3,928 total variants including both novel and ‘hotspot’ targets (i.e., the designed SNP targets of the amplicon panel). Transferring these variants to the chromosome-level assembly (GCF_902806645.1) resulted in 3,685 (93.8%) retained SNPs, including 534 hotspot target SNPs. As novel SNPs identified by TVC are only genotyped in samples with at least one non-reference allele, these loci were not useful here because of incomplete calling across samples and were discarded, leaving only hotspot target SNPs (hereafter referred to as the hotspot dataset). A per-sample missing data filter (GR ≥ 70%) was applied to the hotspot dataset, resulting in the retention of 184 out of 240 offspring samples (76.7%; Figure S2B). Filtering per locus for missing data (GR ≥ 70%) removed 101 SNPs, leaving 407 SNPs in the dataset (Figure S3B). Of these SNPs, 153 were monomorphic and were removed, leaving 254 loci in the filtered hotspot dataset.

### Genotyping concordance in loci shared by HD and LD datasets

Of the 1,257 SNPs in the LD dataset, 26 were also present in the filtered HD data, and these loci were used to compare concordance between the platforms in 183 samples (n = 8 parents; n = 175 offspring). The average per individual genotype concordance (GC) was 97.1 ± 3.6% (Figure 2A) and allelic dosage Pearson correlation was r = 0.940 ± 0.164 (Figure 2B). The average per-locus GC was 97.0 ± 4.0% (Figure 2C; Additional File S4).

**Figure 2.**
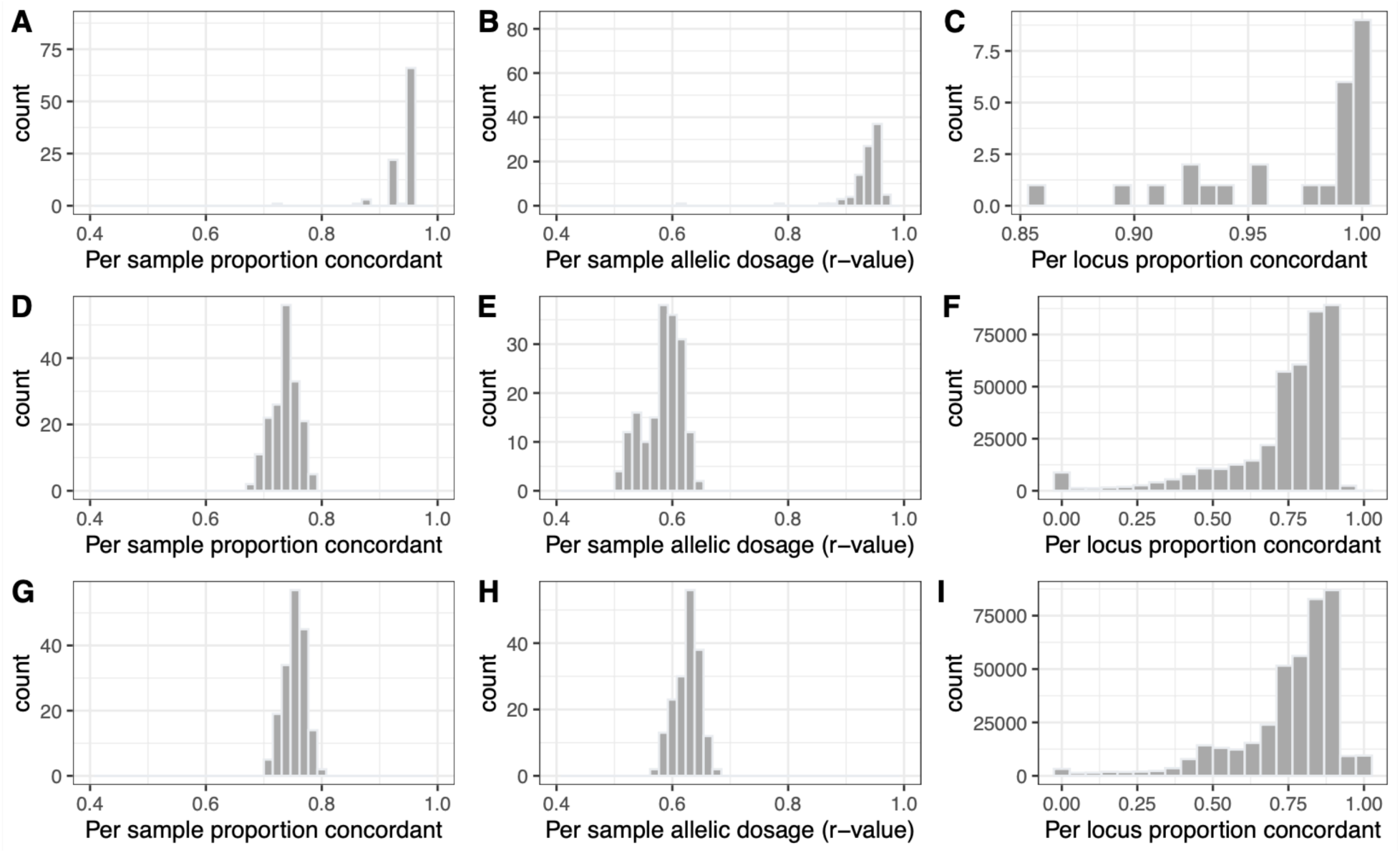
Average per-sample proportions of concordant genotypes (A, D, G), per-sample correlations of allelic dosage (B, E, H), and per-locus concordant genotypes (C, F, I) were determined for loci empirically genotyped independently by both the amplicon panel and whole-genome resequencing (A-C; n = 26 loci and 184 indiv.); for the dataset imputed by AlphaImpute2 (D-F); or by FImpute3 (G-I) relative to empirical genotypes from whole-genome resequencing data (n = 400.9k loci; 176 individuals).

### Mendelian incompatibilities in HD and LD parent-offspring trios

Mendelian incompatibilities (MI) observed from parent-offspring trios indicated that on average (± s.d.), each trio had 4.8 ± 3.4% of the evaluable loci exhibiting MI in the LD panel dataset (n = 240 trios) and 0.3% ± 0.1% in the HD dataset (n = 176 trios; Additional File S5). High MI loci in the panel data included 11 loci exhibiting MI in ≥ 30% of the trios and 254 loci (or 20.2% of total LD panel SNPs) that exhibited MI in ≥ 10% of trios. High MI loci in the HD dataset were less frequent, with only four loci exhibiting MI in ≥ 30% of the trios and 3,765 loci (or 0.89% of HD data SNPs) that exhibited MI in ≥ 10% of trios. The percentage of MI was notably lower for family F117, with an average of 0.06% MI loci per individual (Figure S5).

### Imputation and accuracy evaluation

Of the 424,951 SNPs in the HD+LD dataset, 402,115 were located on chromosomes and were therefore used for imputation. This included 400,896 and 400,889 loci shared with the HD offspring dataset for AlphaImpute2 and FImpute3, respectively, and these were used to evaluate imputation accuracy. For the 176 offspring with HD data, the average (± s.d.) GC per sample between the HD and imputed data was 73.6 ± 2.3% and 75.3 ± 1.9% for AlphaImpute2 and FImpute3, respectively (Table 2; Figure 2) and the average per-sample allelic dosage Pearson correlation was r = 0.584 ± 0.033 and r = 0.626 ± 0.021, respectively (Figure 2).

**Table 2.** Imputation evaluation for all datasets including the standard workflow (‘*WGRS and amplicon panel, full dataset*’) with two generations with imputation by either AlphaImpute2 (AI2) or FImpute3 (FI3). Additionally shown are the three- and two-generation datasets generated *in silico* using the whole-genome resequencing and RADseq data (‘*WGRS + RAD, and in silico panel*’), as well as the WGRS data reduced to be a similar proportion of loci in the HD and LD data as the *in silico* datasets (‘*WGRS and amplicon panel, subset*’). Evaluation metrics included average (± s.d.) per-sample genotype concordance and per-sample allelic dosage (Pearson r-value). The number of imputed loci includes the LD loci, but these were excluded before assessing concordance.

| Dataset | WGRS and amplicon panel, full dataset |  | WGRS + RAD, and *in-silico* panel |  | WGRS and amplicon panel, subset |
| --- | --- | --- | --- | --- | --- |
| *Generations analyzed* | 2 | 2 | 3 | 2 | 2 |
| *Imputation program* | AI2 | FI3 | FI3 | FI3 | FI3 |
| *Low density (LD) panel loci* | 1,257 | 1,257 | 924 | 924 | 990 |
| *Imputed loci* | 402,115 | 402,108 | 4,657 | 4,657 | 4,657 |
| *Per-sample genotype concordance (%)* | 73.6 ± 2.3 % | 75.3 ± 1.9 % | 84.5 ± 3.4 % | 79.3 ± 2.9 % | 76.3 ± 3.0 % |
| *Per-sample allelic dosage (r-value)* | 0.584 ± 0.033 | 0.626 ± 0.021 | 0.804 ± 0.031 | 0.728 ± 0.025 | 0.693 ± 0.027 |

Imputed per-individual genotypes of homozygous reference, heterozygous, and homozygous alternate were on average 341.6k (85%), 27.8k (6.9%), and 32.8k (8.2%), respectively for AlphaImpute2, and 345.0k (85.8%), 28.2k (7.0%), and 28.9k (7.2%), respectively for FImpute3 (Table 1; Additional File S3; Additional File S2). All offspring had similar proportions of each genotype (Additional File S2). Compared with the empirical HD dataset (see above), FImpute3 data had 24% more homozygous reference genotypes, 20% fewer heterozygotes, and 4% more homozygous alternates. The results for AlphaImpute2 were similar (Table 1; Additional File S2). Individual counts and proportions for all individuals can be found in Additional File S3.

Allele frequency differences were also explored between the empirical and imputed datasets by comparing all 400.9k loci and 176 offspring common to both. The empirical dataset had an average (± s.d.) allele frequency of 0.181 ± 0.199 (median = 0.106), whereas the imputed dataset (FImpute3) had an average of 0.115 ± 0.269 (median = 0; Table 3). On average, the empirical minus imputed AF per locus was 0.066 (median = 0.078); the empirical data consistently had higher allele frequencies than the imputed data. This was true for both imputation methods (AlphaImpute2 and FImpute3). Many low AF variants were imputed as fixed homozygous reference, and high AF as fixed homozygous alternate (Figure 3).

**Figure 3.**
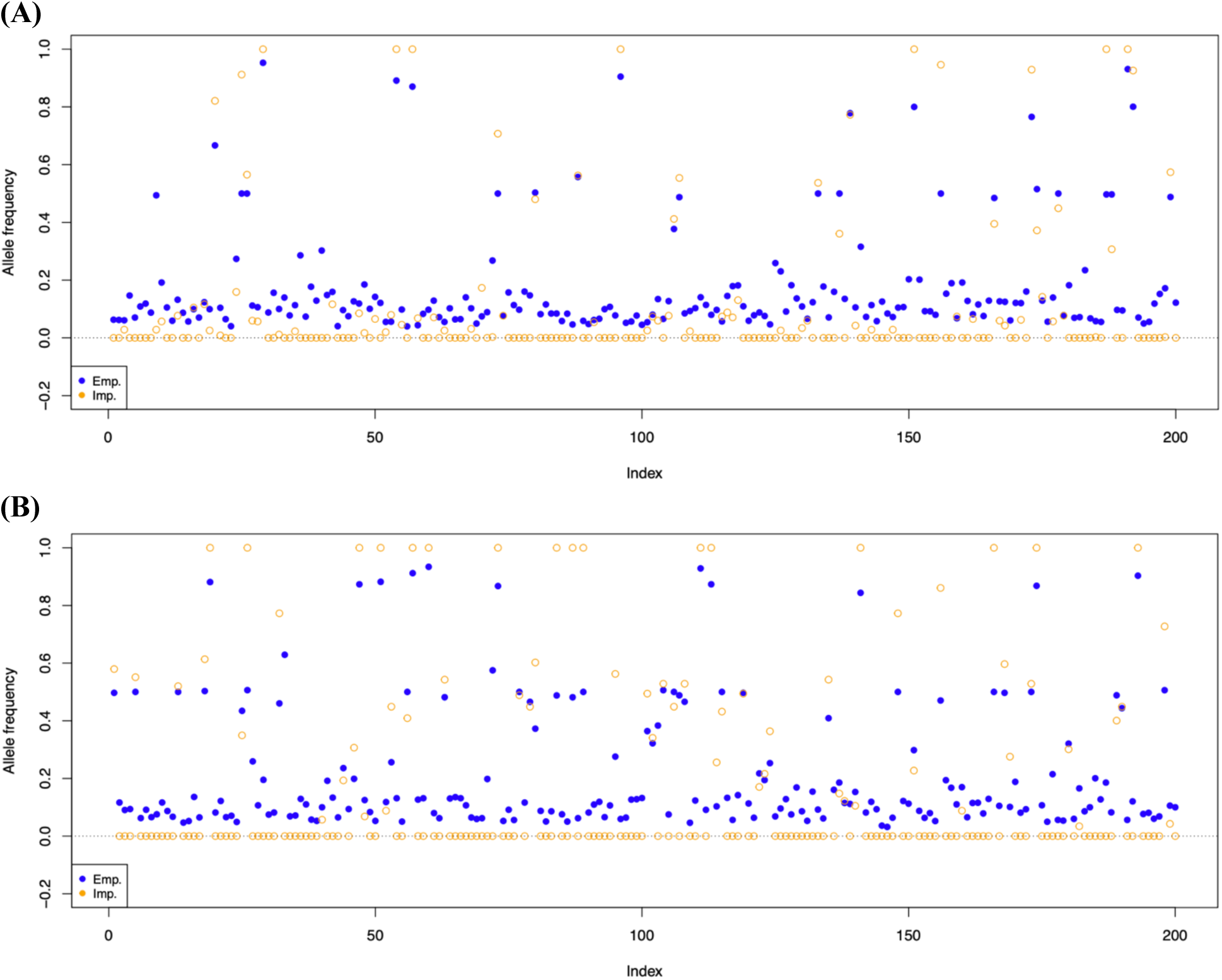
Randomly selected loci (n = 200) shown with allele frequency from both empirical (blue, closed circle) and imputed data (orange, open circle) for (A) AlphaImpute2; and (B) FImpute3 indexed along the x-axis.

**Table 3.** The average, standard deviation (s.d.), and median allele frequencies of 400.9k loci in the empirical, AlphaImpute2 (AI2) or FImpute3 (FI3) imputed datasets, as well as the per-locus AF difference (empirical minus imputed).

|  | **Empirical**<br>**AF** | **Imputed**<br>**(AI2) AF** | **Empirical minus**<br>**imputed (AI2) AF** | **Imputed**<br>**(FI3) AF** | **Empirical minus**<br>**imputed (FI3) AF** |
| --- | --- | --- | --- | --- | --- |
| *Average* | 0.181 | 0.107 | 0.074 | 0.115 | 0.066 |
| *s.d.* | 0.199 | 0.249 | 0.105 | 0.269 | 0.138 |
| *Median* | 0.106 | 0 | 0.073 | 0 | 0.078 |

### Grandparent genotyping, in silico panel development, and imputation evaluation

Published grandparent RADseq sequence data (Divilov et al. 2023b) were genotyped using an approach similar to that used for whole-genome resequencing data (see Methods), producing 128,300 quality-filtered SNPs, of which 4,976 loci were shared with the whole-genome resequencing dataset. The 4,976 loci were used to create an *in silico* LD panel of 1,000 loci present in all three generations (grandparents, parents, and offspring), and all 4,976 loci were kept for grandparents and parents. After removal of non-chromosomal loci and the loci present in the *in silico* panel, 3,733 imputed loci were retained to evaluate accuracy.

Empirical and imputed concordance evaluation for the three-generation *in silico* dataset provided a GC of 84.5 ± 3.4% with an allelic dosage correlation Pearson r = 0.804 ± 0.03. This was approximately 10% higher in GC and a 0.18 higher r-value than the best results with the two-generation amplicon panel dataset (Table 2). These values were reduced to GC = 79.3 ± 2.9% and r = 0.728 ± 0.03 when only the two generations (parents and offspring) were included in the *in silico* dataset, which was still significantly higher than the amplicon panel dataset (Table 2). However, the down-sampled empirical amplicon panel dataset with a similar ratio of LD and HD loci to the ratio in the *in silico* analysis yielded a GC of 76.3 ± 3.0% and r = 0.693 ± 0.03, which is very similar to the *in silico* dataset with two generations and slightly higher in correlation r-value (+0.07) than the full empirical amplicon panel dataset (Table 2). Therefore, the three-generation *in silico* dataset provided the best results, and the two-generation *in silico* dataset was similar to the two-generation amplicon panel datasets; however, there was a slight advantage (as observed by higher allelic dosage but not GC) when the LD panel was more similar in number to the number of imputed loci.

### GWAS analyses for OsHV-1 survival

Of the 240 samples and four families, families F114 and F115 had low mortality rates (13.3% and 17.5%, respectively) compared to F116 and F117 (64.9% and 51.7%, respectively; Table 4; Figure 4). Of the 240 offspring, six did not have a recorded survival phenotype at the end of the experiment and were removed (n = 234 offspring retained). Only 176 samples were available in the whole-genome resequencing (HD) dataset, with only 28 of the 86 mortality samples having high quality data. Given the high mortality drop out in the HD dataset, the number of mortalities per-family was not expected to provide sufficient power for a GWAS analysis. To address this sample size challenge, the amplicon panel-based imputation was explored to recover as many samples as possible (see workflow in Figure 1).

**Figure 4.**
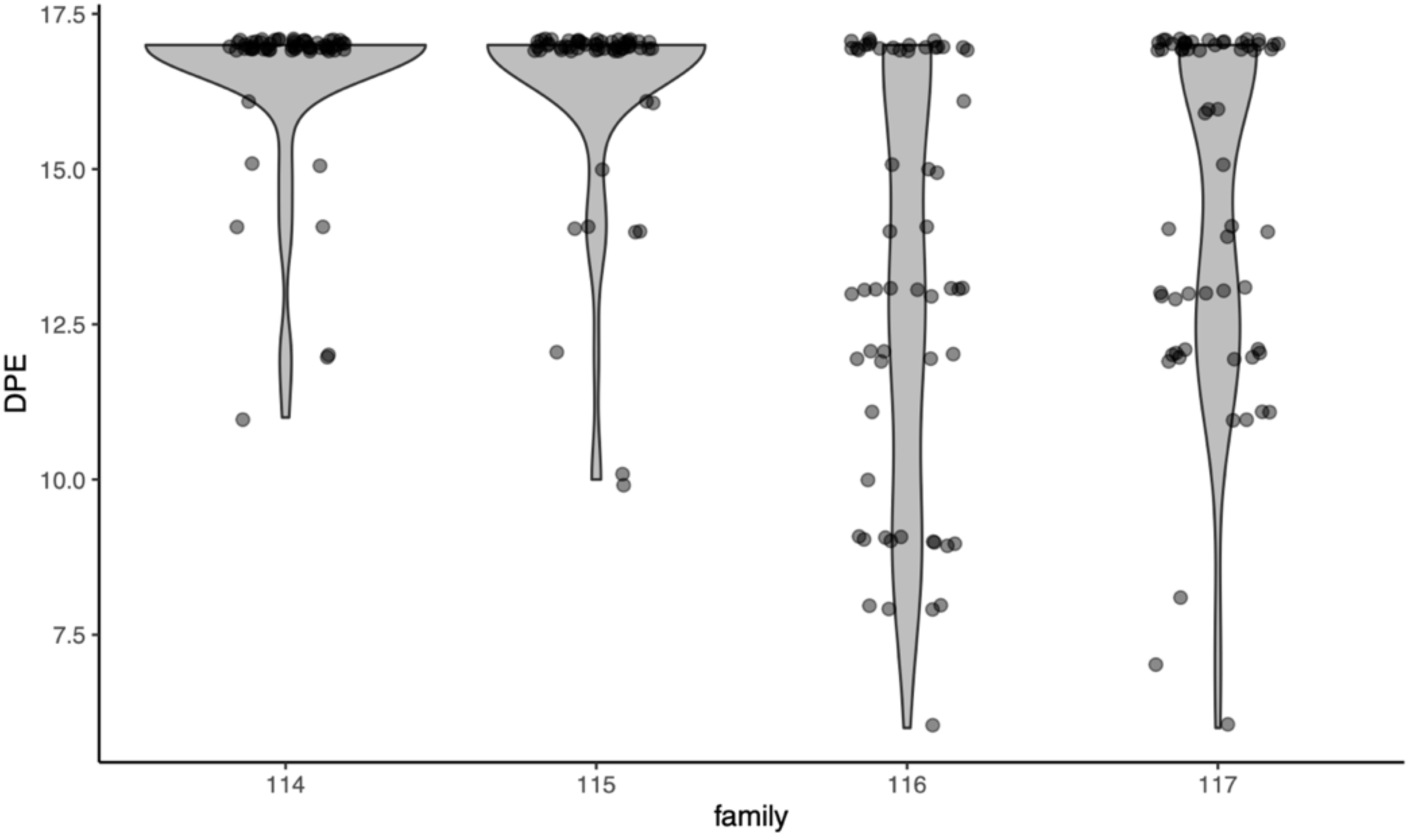
Per-family mortality expressed as days post-exposure (DPE) following the OsHV-1 exposure in each of the four families (n ∼ 60 offspring per family). Survivors were assigned a DPE value of 17 (i.e., the day after the trial concluded).

**Table 4.** Sample sizes and percentage of total available samples retained per family and survival in the OsHV-1 trial (M = mortality; S = survivor) based on all samples and those analysed in four datasets: whole-genome resequencing data; amplicon panel data using target hotspots only; amplicon panel using *de novo* identified SNPs; and imputed data using high-density parents and the *de novo* SNPs in the low-density panel. The amplicon panel datasets were filtered at a per-individual genotyping rate of 70%. Low retention of samples in the WGRS data was due to poor DNA quality of extracted samples.

| **Dataset** | **F114** |  | **F115** |  | **F116** |  | **F117** |  | **TOTAL** |
| --- | --- | --- | --- | --- | --- | --- | --- | --- | --- |
|  | **M** | **S** | **M** | **S** | **M** | **S** | **M** | **S** |  |
| *All samples* | 8 | 52 | 10 | 47 | 37 | 20 | 31 | 29 | 234 |
| *WGRS only* | 4<br>(50%) | 52<br>(100%) | 2<br>(20%) | 47<br>(100%) | 11<br>(29.7%) | 20<br>(100%) | 11<br>(35.5%) | 29<br>(100%) | 176<br>(75.2%) |
| *Amp. panel, hotspots* | 3<br>(37.5%) | 51<br>(98.1%) | 5<br>(50%) | 47<br>(100%) | 28<br>(75.7%) | 19<br>(95%) | 12<br>(38.7%) | 19<br>(65.5%) | 184<br>(78.6%) |
| *Amp. panel, de novo* | 8<br>(100%) | 52<br>(100%) | 9<br>(90%) | 47<br>(100%) | 34<br>(91.9%) | 20<br>(100%) | 22<br>(71%) | 28<br>(96.6%) | 220<br>(94.0%) |
| *Imputed* | 8<br>(100%) | 52<br>(100%) | 10<br>(100%) | 47<br>(100%) | 37<br>(100%) | 20<br>(100%) | 31<br>(100%) | 29<br>(100%) | 234<br>(100%) |

The *de novo* panel loci dataset included 220 individuals (n = 73 mortalities and 147 survivors), and the imputed datasets included all 234 samples with phenotypes (n = 86 mortalities and 148 survivors). GEMMA analyzed 1,117 of the 1,257 *de novo* panel SNPs, 58,556 of the 402,115 SNPs (14.6%) imputed by AlphaImpute2 and 73,984 of the 402,108 SNPs (18.4%) imputed by FImpute3. One locus passed significance in the panel-only dataset, at 39.65 Mbp on Chr8 (Figure 5A; Figure 6A; Additional File S6). The GWAS with AlphaImpute2-imputed loci found several significant association peaks (10 SNPs), all located on Chr8. These SNPs were at 12.48 Mbp, 16.25-16.31 Mbp, 35.92 Mbp, and 36.05 Mbp (Figure 5B; Figure 6B). The GWAS with FImpute3-imputed loci also found several significant peaks (86 SNPs), at 15.84 Mbp (1 SNP), 31.84-31.87 Mbp (6 SNPs), 35.48 Mbp (3 SNPs), and 42.15-42.79 Mbp of Chr8 (76 SNPs; Figure 5C; Figure 6C). Notably, no significant SNPs were shared between the GWAS results using the AlphaImpute2 and FImpute3 imputation datasets. There was no detected association at the previously-associated Chr8 survivorship locus location (9.72 Mbp), with the closest being the association at 12.48 Mbp of Chr8 in the AlphaImpute2 GWAS. Predicted genes within the association regions from the *de novo* panel only, AlphaImpute2, and FImpute3 analyses are shown in Additional File S7. The predicted gene with the most direct potential link to survivorship based on antiviral properties is *interleukin-12 receptor subunit beta-2* at 42.69 Mbp of Chr8, but there were many other predicted annotations within the windows, given the number of associated regions. The imputation by AlphaImpute2 did not provide usable loci for Chr1, as these were all removed by GEMMA during the analysis (Figure 5B), and this chromosome has a comparatively small number of loci in the amplicon panel.

**Figure 5.**
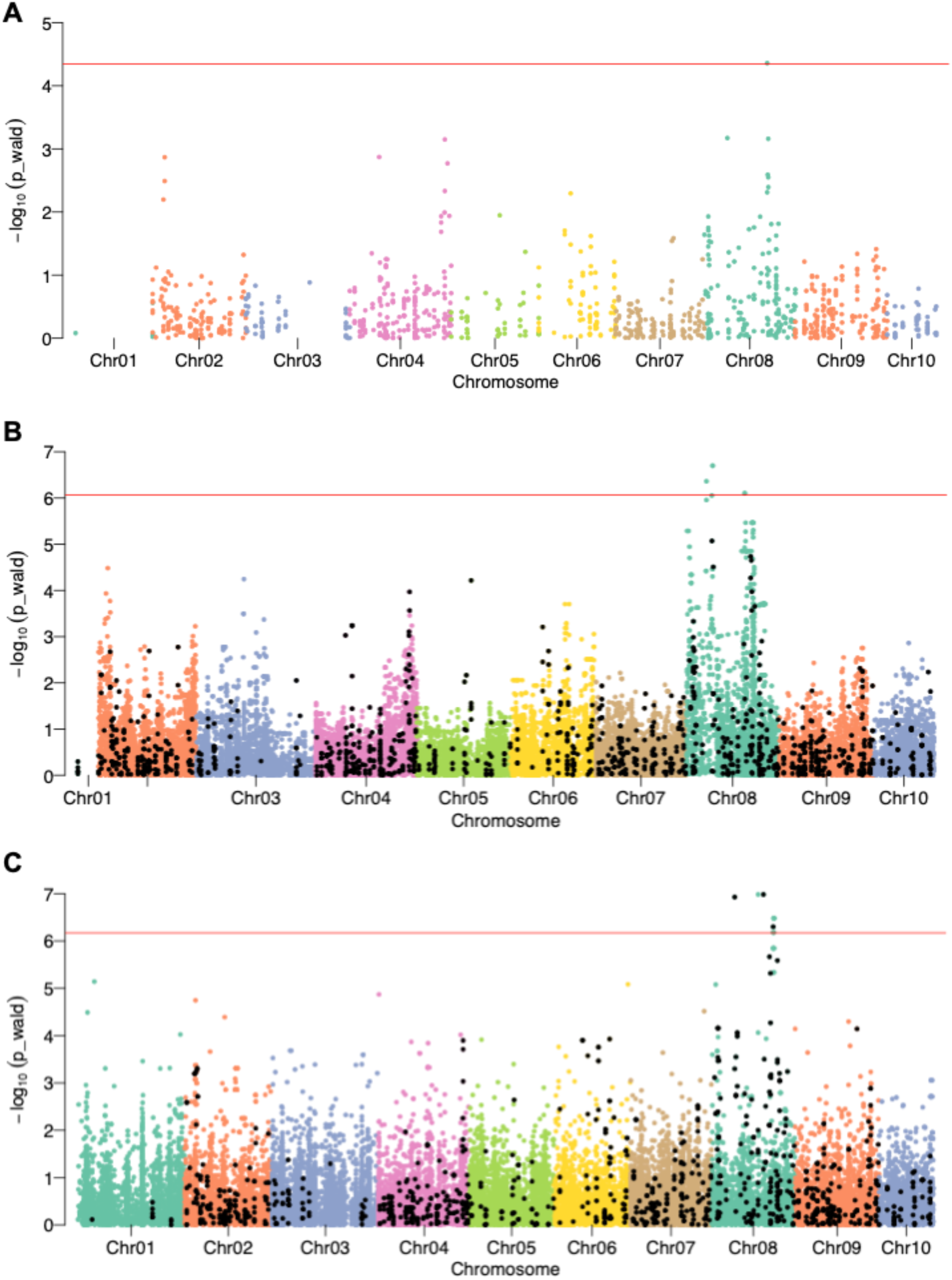
Manhattan plots for survival after exposure to OsHV-1 in (A) amplicon panel *de novo* SNPs (n = 1,117 loci and 214 individuals); (B) imputed data with AlphaImpute2 (n = 58,556 loci and 234 individuals); and (C) imputed data with FImpute3 (n = 73,984 loci and 234 individuals). The red horizontal line represents the genome-wide Bonferroni-corrected p-value significance threshold of 0.05.

**Figure 6.**
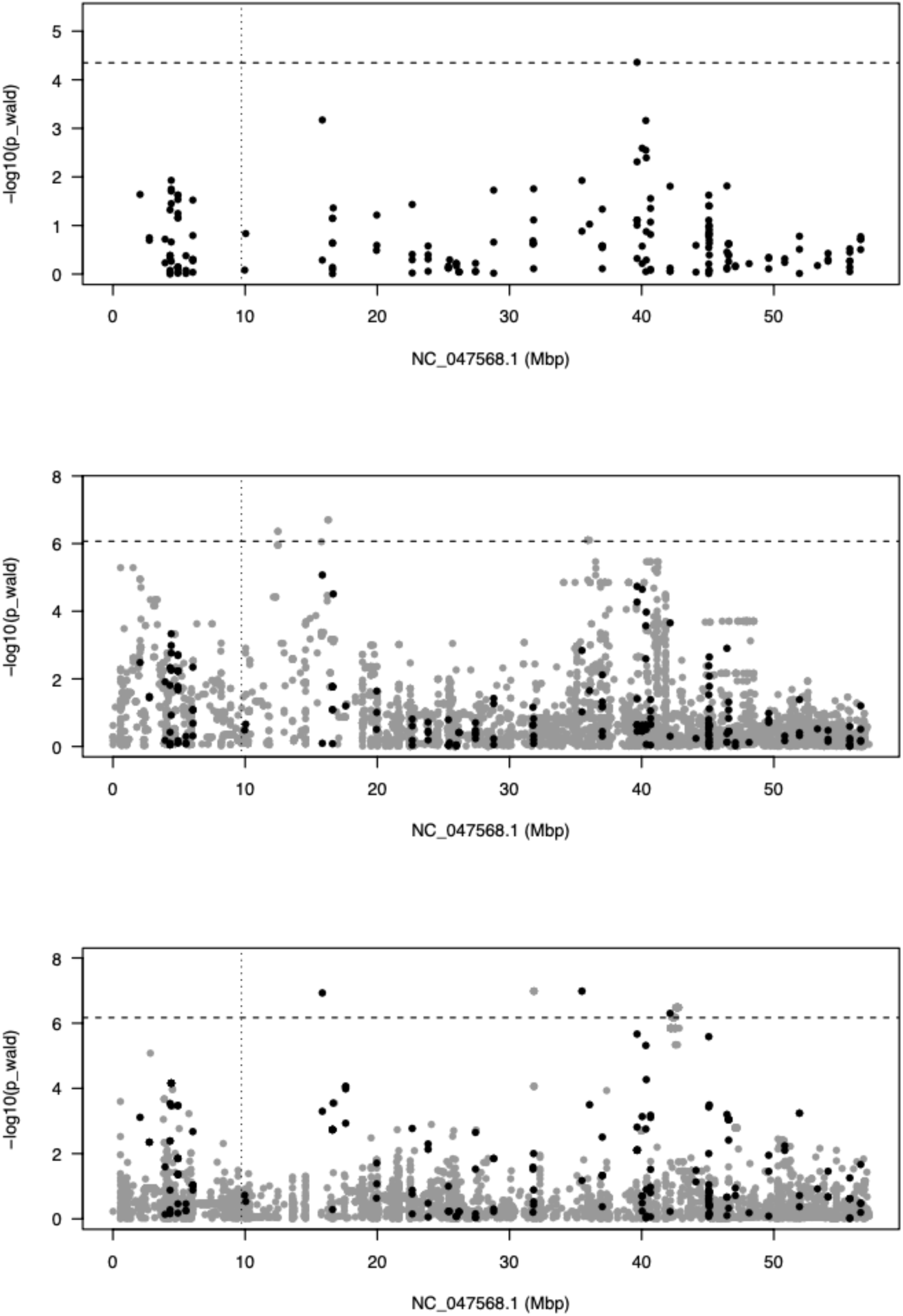
Manhattan plot for associations to survival after exposure to OsHV-1 for the target chromosome NC_047568.1 only (i.e., Chr8), shown with results from (A) amplicon panel *de novo* SNPs; (B) imputed data by AlphaImpute2; and (C) imputed data by FImpute3. The horizontal hatched line is the Bonferroni-corrected significance cutoff, and the vertical hatched line is the position of the previously-identified Chr8 survival locus. Black dots are those from the amplicon panel, and grey are imputed loci.

## Discussion

Here we used a low-density Pacific oyster amplicon panel for genotyping four families of selectively bred oysters with directly measured OsHV-1 survival phenotypes from a laboratory exposure trial. We genotyped the parents using whole-genome resequencing (WGRS) and imputed offspring genotypes for all 10 chromosomes. We evaluated imputation accuracy by comparing offspring genotypes to a hold-out set of WGRS offspring genotypes that were not used for imputation. We then used the imputed genotypes in a genome-wide association study (GWAS) for OsHV-1 survival to improve our understanding of the genetic architecture of the survival trait. Below we discuss (1) details of imputation accuracies; (2) the utility for GWAS and new insights on the genetic architecture of OsHV-1 survival in the selectively bred families; and (3) the potential for these results to guide future studies in GS in Pacific oysters, considered to be the next frontier for genetic improvement in agricultural and aquaculture species (Meuwissen et al. 2016; Voss-Fels et al. 2019; Houston et al. 2020; Yáñez et al. 2022; Andersen et al. 2025).

### Accuracy and characterization of imputed data

Imputation accuracy was influenced by the number of generations present in the dataset. This was demonstrated by comparing the accuracy in the two-generation (i.e., parents, offspring) analysis and in the three-generation (i.e., grandparents, parents, offspring) *in silico* analysis. Because we only had two generations of data empirically genotyped with the amplicon panel, we were not able to achieve the maximum imputation accuracy that would be expected had we included the grandparent data at high density and with the amplicon panel. With the amplicon panel data, the best imputation accuracy was obtained with FImpute3 (GC = 75.3% and r = 0.626). In contrast, the three-generation *in silico* panel imputation had significantly higher imputation accuracy (GC = 84.5% and r = 0.804). This was expected, as *in silico* studies indicate a minimum requirement of two parental generations genotyped at high density (HD) for maximal imputation accuracy (Delomas et al. 2023, 2024). Higher haplotype resolution can be obtained from deeper pedigrees (Fraslin et al. 2023). An additional detail observed here was that when the amplicon panel two-generation data was reduced to a similar ratio of imputed loci to low-density (LD) panel loci (i.e., 3,733 imputed loci to 990 panel loci), the allelic dosage correlation was improved (r = 0.693) relative to the full dataset (i.e., 424,925 imputed loci to 1,257 panel loci; r = 0.626). Given that the results were affected by both the number of generations genotyped at HD, and the ratio of HD:LD loci, although we expect substantially better results with three generations included, concordance may not be as high as that observed in the *in silico* panel dataset here because of the larger HD to LD ratio that would occur. The *in silico* dataset has the additional benefit of using similar sequencing technology for both the HD and LD data as opposed to expected standard applications that would use different library preparation methods and sequencing platforms between the amplicon panel and short read sequencing.

Imputation accuracy has been previously investigated in Pacific oysters *in silico* (Delomas et al. 2023; Kriaridou et al. 2023; Delomas et al. 2024), and the empirical application here brought new insights. First, we observed that it was necessary to use *de novo* genotyping for the amplicon panel (as per Thompson et al. 2025) instead of only the default target SNPs to obtain a sufficient number of high-quality genotypes and retained samples after quality filters. The *de novo* dataset retained 214 of 234 offspring with 1,257 SNPs, whereas the default target approach would have only retained 184 offspring and 254 SNPs. The value of *de novo* SNP identification agrees with the conclusions of Delomas et al. (2024). Similar to Thompson et al. (2025), *de novo* genotyping was found to be particularly valuable when applying the amplicon panel to populations other than those included in the panel design, including populations that have been subject to extensive selective breeding. This is promising, as it demonstrates that the panel can be used broadly and will likely have greater longevity than if the target variants were the sole focus.

Another insight from this empirical study was the effect of using a different platform for the LD and HD genotypes. Even though the few loci that were shared between the platforms had high concordance, there was a significantly higher proportion of loci exhibiting Mendelian incompatibilities (MI) in the amplicon panel data than in the WGRS data (i.e., a 20x higher proportion of loci exhibited MI in at least 10% of inspected trios in the panel). Based on known challenges with null alleles when amplifying DNA from Pacific oyster (Hedgecock et al. 2004), the higher MI occurrence in the amplicon panel data may be due to a higher prevalence of null alleles in the amplicon panel data than in the WGRS data. Putative null alleles have been previously identified in the amplicon panel (Sutherland et al. 2024; Thompson et al. 2025). The null alleles were determined to be largely population-specific, and were demonstrated to improve parentage assignments once removed (Sutherland et al. 2024; Thompson et al. 2025). Null alleles also occur in SNP arrays for the eastern oyster (Guo et al. 2023; Xuereb et al. 2023), and may be caused by mutations in primer sites (Hedgecock et al. 2004) and/or structural variations such as small and large deletions (Guo et al. 2023). MI may also be caused by *de novo* mutations, or other forms of genotyping error, such as false homozygotes in low-quality samples (Xuereb et al. 2023). Guo et al. (2023) suggested that the main cause of null alleles in an eastern oyster SNP chip was from deletions (structural variants), in part because clusters of null alleles were observed in the genome that were flanked by loci exhibiting regular inheritance. In the present study, it is possible that the high MI rate was due to the prevalence of null alleles caused by disrupted primer sites (as discussed in Thompson et al. 2025), since the WGRS data did not show similarly high rates of MI. If deletions are causing null alleles in the present study, it is also possible that the WGRS variant calling is better able to detect, and appropriately deal with them. This higher sensitivity would likely filter out these regions (i.e., deletions) before MI detection. These empirical observations will help guide future studies that depend on amplicon panel-based imputation.

Imputation inaccuracies and challenges that have been observed to occur in Pacific oysters (Kriaridou et al. 2023) may be related to various characteristics of the species, including high polymorphism. Typically, high SNP polymorphism would be expected to be beneficial for imputation; high SNP density can be used to resolve haplotypes across genomic regions and be informative for determining identical-by-descent segments (Abney and ElSherbiny 2019). If high polymorphism levels also characterize numerous rare variants (i.e., low MAF SNPs), this can challenge imputation as low MAF SNPs are more difficult to accurately impute (Blue et al. 2014; Ros-Freixedes et al. 2020; Si et al. 2021). If imputation program models applied expect and use Mendelian ratios (Wijsman 2016), and if these do not occur due to null alleles or genotype-dependent offspring mortality, this could also challenge imputation. Pacific oyster families deviate from expected genotype ratios during development, at least in part, due to larval mortalities (Durland et al. 2021) that are due to high genetic load and negative allelic combinations (Plough 2016). In the present study, empirical genotypes followed expected modes of MAF that would occur from a biparental cross; however, there was also substantial per-locus variation around the modes. Regions with repeat sequences or structural variants also can influence imputation errors (Blue et al. 2014), of note given the extensive genome-wide structural variation known to occur in oysters (Modak et al. 2021; Jung et al. 2025). Regardless of the cause, lower imputation accuracy in Pacific oysters than in finfish is expected based on comparative *in silico* analyses of multiple species (Kriaridou et al. 2023). The accuracy of genotype calls is the main goal of imputation; in the present study, we observed clear and systematic skews in genotype proportions between the imputed and empirical datasets, with significant enrichment for homozygotes and depletion of heterozygotes in the imputed data, alongside a systematic reduction in allele frequency in the imputed data. The accuracy presented here in the two-generation, amplicon-panel-based dataset (r = 0.626) was lower than that obtained by Kriaridou et al. (2023) in a two-generation, *in silico* dataset (r = 0.775), and Delomas et al. (2023) in a two-generation, *in silico* dataset with 250 or 2,000 loci in the LD panel (r = 0.7 and 0.85, respectively). It is also worth noting that strikingly high concordance has been presented for low-coverage (∼2.8x) whole-genome resequencing for Pacific oysters imputed without a reference panel (GC = 93.6% and R^2^ = 0.860; Yang et al. 2024).

We expect that future studies using at least three generations will improve accuracy, but it is worth considering whether the results generated here are sufficient for downstream applications, especially considering cost savings with the present approach relative to other approaches. Although per-sample costs are expected to vary and may change over time, the general averages that are available for the species point to major savings that can be obtained by using the amplicon panel combined with imputation, considering amplicon panel genotyping costs at $15 USD, RAD-seq at $35 USD, SNP chip genotyping at $60 USD, and whole-genome resequencing (10X coverage) at $90 USD (also see Meek and Larson 2019). As an example, genotyping 240 individuals would cost $3,600 by amplicon sequencing, $8,400 by RAD-sequencing, $14,400 by SNP chip, or $21,600 by WGRS. To conduct imputation, additional high-density sequencing for imputation purposes would need to be added to these costs, for example, for parents and grandparents by WGRS.

### Applying amplicon panel-based imputation in a GWAS for OsHV-1 resistance

Using amplicon panel-based imputation, we were able to recover a significant number of samples that could not be genotyped using WGRS due to poor quality DNA, including many of the mortalities from the OsHV-1 laboratory challenge. Shellfish are expected to undergo rapid DNA degradation post-mortem, and genomic DNA extraction from oysters can be challenging (Adema 2021). As a result, 64 of the 240 offspring samples could not be genotyped using WGRS, significantly reducing the sample size for GWAS. As the amplicon panel was less sensitive to degraded DNA, *de novo* genotyping and imputation were conducted. This approach identified significant associations with OsHV-1 survivorship on the expected chromosome for these families (i.e., Chr8; Divilov et al. 2023b, 2023a). However, the observed peaks were not located near the previously reported survivorship-associated SNP (9.72 Mbp; Divilov et al. 2023b), or the other associated regions identified using allele frequency studies through a field exposure to OsHV-1 (Divilov et al. 2023a), all of which were identified in the first 10 Mbp of Chr8. Therefore, in agreement with Divilov et al. (2023a), we find further associations on Chr8 but consider the resistance mechanisms of these families to involve a wider variety of regions and putative genes.

The exact locations of the associations observed here varied slightly between datasets depending on the imputation software applied, but both imputation datasets identified peaks near 35-36 Mbp of Chr8 (AlphaImpute2 at 35.9 Mbp; FImpute3 at 35.5 Mbp). The most accurate imputation approach evaluated here, FImpute3, found the most loci associated with survivorship to OsHV-1 trial between 42.15-42.79 Mbp of Chr8 (n = 76 SNPs). Although this region contains many predicted genes, a gene of interest within the associated regions with known antiviral activity is *interleukin-12 (IL-12) receptor subunit beta* at 42.69 Mbp. In vertebrates, IL-12 regulates the adaptive and innate immune responses and is involved in antiviral response (Trinchieri 2003), and has been shown to have potent antiviral activity against herpesvirus (Carr et al. 1997). Cytokines are known to be induced by pathogen stimulation in Pacific oyster haemocytes (Zhang et al. 2023), and an IL-12 homolog is present in Pacific oyster with an antibacterial role (Xin et al. 2021). In vertebrates, IL-12 induces the production of *interferon gamma* (IFN-**γ**) (Trinchieri 2003), which in turn activates the interferon response factor 1 (IRF1) pathway (Tomita et al. 2003). Interestingly, basal overexpression of IRF2 has previously been documented in relation to the protective allele of the Chr8 marker (Divilov et al. 2023b). IRF1 and IRF2 function as transcription factors with distinct but complementary roles in cellular responses to viral infection (Ren et al. 2015). Due to the discrepancies between the two imputed datasets, and the differences in associated regions relative to earlier studies, further investigation would be required to identify causative locI conferring increased survival in field and laboratory challenge environments. Chr8 appears to have multiple relevant regions associated with OsHV-1 laboratory challenge survivorship, although the exact mechanisms remain elusive.

In addition to the lack of signal in the present GWAS at the 9.72 Mbp locus, other uncertainty about the predictability of the Chr8:9.72 Mbp OsHV-1 survivorship marker is notable from the present study. Specifically, all four families in the exposure trial were expected to have similar genotype frequencies at the locus (all parents being heterozygous for the SNP), but significant variation in mortality was observed between the families in the trial. One possible explanation is that the genotype frequencies of the 9.72 Mbp marker could have been skewed prior to the infection trial, resulting in basal differences in frequencies before OsHV-1 challenge. Since the marker was not genotyped here, as it is not present in the current version of the amplicon panel, we do not have information about the genotypes at this marker before the trial started. Two of the four families (F114 and F115) had very low mortality rates, which differed from F116 and F117. The low mortality reduced GWAS detection power and raises the question of whether the Chr8:9.72 Mbp marker is still predictive of OsHV-1 survival in these families. Another possible explanation for this discrepancy between families is that the Chr8:9.72 Mbp locus was identified in relation to field survival (Divilov et al. 2023b), whereas the present study used survival data from a laboratory OsHV-1 infection trial. Field survival could be influenced by additional factors other than only OsHV-1 resistance or tolerance. However, the Chr8:9.72 Mbp marker alternate allele was associated with survival in the same infection trial as the present study in different families with offspring that were either all homozygous reference or all heterozygous for the locus (Lunda et al. *in review*), which provides evidence that the locus is directly associated with OsHV-1 survival. It is also possible that the detection of associated genomic regions in the present study could be related to read mapping or imputation inaccuracies, but this still would not explain the two families with higher-than-expected survival rates. Furthermore, the distal region of the chromosome was also identified using the *de novo* panel loci only.

Investigation of the genetic underpinnings of this trait will need to be continued in future work. Unpublished data from our research group suggests that the linkage between the putative causative major effect locus and the marker applied in MAS may have broken down for at least some of the families in this cohort, which could result in a loss of signal at Chr8:9.72 Mbp as observed here. We are currently investigating this phenomenon. Similar to the conclusions of Divilov et al. (2023a), we suspect that the observed peaks in the present study represent other significantly associated large effect loci on Chr8. This would suggest that the OsHV-1 survival phenotype has at least oligogenic architecture on Chr8 in these families. The consistent identification of different regions of the same chromosome make this an interesting area of study in relation to resistance or tolerance to viral infection in an invertebrate. Furthermore, the breakdown of linkage between the marker used for MAS and the causative marker involved in OsHV-1 survival highlights the limitation of using linked markers in selective breeding of oysters, as the predictive utility of a linked marker may be reduced or lost relatively quickly.

### Applying amplicon panel-based imputation in future genomic selection approaches

Although the application of GWAS was of interest in the current study, the main proposed applications of imputation to support breeding are for genomic selection (GS) (e.g., Delomas et al. 2023; Kriaridou et al. 2023; Delomas et al. 2024). Therefore, it is relevant to discuss whether imputation accuracy obtained in the present study, as described above, is sufficiently reliable for GS. A recent study on rainbow trout *Oncorhynchus mykiss* demonstrated that even with lower imputation accuracy (e.g., r = 0.58 for a 300 SNP LD panel) genomic prediction accuracy did not strongly deviate from that obtained with higher accuracy imputation (Fraslin et al. 2023). Whether this same conclusion would hold in empirical studies using the highly polymorphic Pacific oyster is not yet known and will be important to empirically investigate in future work. However, given that we exceeded this genotype concordance with the two-generation imputation here (i.e., r = 0.626), and that this would be relatively easily improved with the addition of a third generation genotyped at HD, these results are promising for the application of the amplicon panel-based imputation for GS. The same level of imputation accuracy (i.e., r = 0.626) was observed by Fraslin et al. (2023) using between 300-500 SNPs in rainbow trout (two generations), and in the simulations by Delomas et al. (2023) using approximately 100 loci and only two generations.

Varying degrees of imputation accuracy have resulted in robust GS results in Pacific oyster simulations (Delomas et al. 2023, 2024). Fraslin et al. (2023) attributed the robustness of genomic prediction, even in the presence of imputation inaccuracies, to the strong linkage disequilibrium that occurs through the use of GS on close relatives (e.g., full-sibs), which is typical in aquaculture studies. This results in increased short- and long-range linkages and large shared haplotype blocks. It would be worthwhile to consider recombination rate variation in Pacific oysters to evaluate how large these shared haplotype blocks are across siblings. Yang et al. (2024) also found the highest imputation accuracy in aquaculture lines rather than outbred wild lines, which was attributed to increased LD from family structure. From their study that demonstrates high effectiveness for GS with low marker numbers, Fraslin et al. (2023) propose using 300-500 SNPs in low-cost amplicon panels combined with imputation from HD relatives, which is what we have conducted here. The use of amplicon panels in this manner brings the additional benefit of being able to confirm pedigrees through parentage analysis using the genotypes required for GS (Fraslin et al. 2023; Sutherland et al. 2024), or even to add putatively functional loci to enable MAS alongside GS.

## Conclusions

This study is the first empirical demonstration of genome-wide imputation in Pacific oysters using a low-density amplicon panel combined with whole-genome resequencing data. These results are expected to guide future work in shellfish genomic selection. The cost of whole-genome resequencing of many individuals, even at low coverage, is prohibitive for most operations. Aquaculture focuses on a wide variety of species, many of which are produced in regionally distinct environments, and economies of scale are sparse in aquaculture. As demonstrated here, the amplicon panel and imputation approach can be used for genotyping and expanded to a genome-wide scale. This should enable small operations to use advanced GWAS and GS methods and large operations to expand genotyping coverage across their programs, although additional empirical demonstration is required for GS. In this work, we provided tissue to a sequencing facility for DNA extraction and sequencing, who in turn provided raw sequence outputs; therefore, very limited in-house laboratory techniques are needed to accomplish genotyping. Improvements in accuracies observed here are expected, particularly through the inclusion of additional parental generations to deepen the pedigree. The varied phenotypic responses to disease of the families studied here, combined with the identification of a different associated genomic regions on Chr8, suggests a complex genetic architecture for OsHV-1 survival, and further studies are needed to understand the drivers of this important phenotype.

## Supporting information

Supplemental Results

## Acknowledgements

Thanks to the staff at the Center for Aquaculture Technologies (CAT) in San Diego and Prince Edward Island for providing information regarding the OsHV-1 challenge. The U.S. Department of Agriculture (USDA) is an equal opportunity provider and employer. Mention of trade names or commercial products in this publication is solely for the purpose of providing specific information and does not imply recommendation or endorsements by the USDA. Thanks to the Editor and two anonymous reviewers for comments on an earlier version of this manuscript.

## Data Availability

The following repositories support this study: Whole-genome resequencing workflow: https://github.com/bensutherland/wgrs_workflow Imputation workflow: https://github.com/bensutherland/impute_workflow Project-specific scripts and instructions: https://github.com/bensutherland/ms_cgig_chr8_oshv1 Genotype files and associated metadata is available on FigShare: 10.6084/m9.figshare.28095440 Raw amplicon panel sequence data and whole-genome resequencing data for the parents and offspring are available under the NCBI SRA BioProject PRJNA1219245. Raw sequencing data for the grandparents were previously published and is available under the NCBI SRA accession number PRJNA873124, including the dam and sire for families 30.055 (SRR21192124 and SRR21192123), 30.058 (SRR21192118 and SRR21192117), 30.065 (SRR21192209 and SRR21192208), and 30.079 (SRR21192111 and SRR21192110), respectively.

## Additional Files

Additional Figures and Tables are provided in the Supplemental Results.

Additional File S1. Per-sample number of reads, alignments, alignment rate, and duplicate level for whole-genome resequencing data.

Additional File S2. Per-sample proportions of genotypes homozygous reference (0/0), heterozygous (0/1), and homozygous alternate (1/1), or missing data in (A) HD data (empirical whole-genome resequencing data) for parents (n = 8) and available offspring (n = 176); (B) imputed data by AlphaImpute2; or (C) imputed by FImpute3 (imputed includes eight parents and all 240 offspring).

Additional File S3. Individual genotype counts for samples present in both the empirical and imputed data.

Additional File S4. Per locus concordance between platforms (panel vs. whole-genome resequencing) or empirical and imputed (both AI2 and FI3).

Additional File S5. Per-sample and per-locus Mendelian incompatibility counts for trios in the LD data (amplicon panel) or HD data (whole-genome resequencing).

Additional File S6. Output of GEMMA analyses for the AlphaImpute2, FImpute3, and *de novo* amplicon panel analyses.

Additional File S7. Predicted gene annotations from OsHV-1 survivorship-associated regions.

