## Supplemental Results for "Pedigree-based genome-wide imputation using a low-density amplicon panel for the highly polymorphic Pacific oyster *Crassostrea* (*Magallana*) *gigas*"

**Table of Contents:**


| | |
|---|---|
| **Figure S1. Allele frequencies per family for expected and unexpected polymorphisms** | **Page 2** |
| **Figure S2. Per-locus percent missing genotypes** | **Page 3** |
| **Figure S3. Per-individual genotyping rate** | **Page 4** |
| **Figure S4. Sample PCA using amplicon panel genotypes** | **Page 5** |
| **Figure S5. Per-individual percentage of loci exhibiting Mendelian incompatibilities** | **Page 6** |
| **Table S1. Variants retained through filtering for whole-genome resequencing and amplicon panel data** | **Page 7** |
| **Table S2. Per-family percentage of loci polymorphic, and unexpected polymorphism** | **Page 8** |

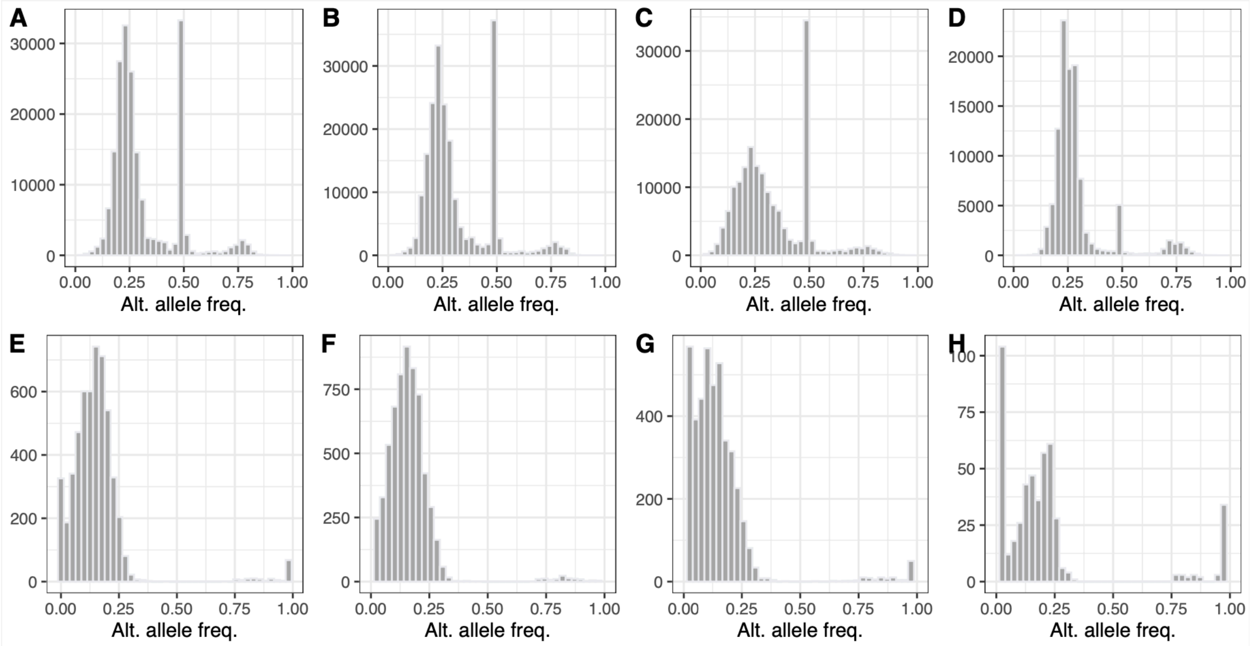


**Figure S1.** Alternate allele frequency for fully genotyped loci in parents that are (A-D) polymorphic in both the parents and in offspring or that are (E-H) monomorphic in parents but polymorphic in offspring. Panels A and E: F114; B and F: F115; C and G: F116; and D and H: F117.

(A)

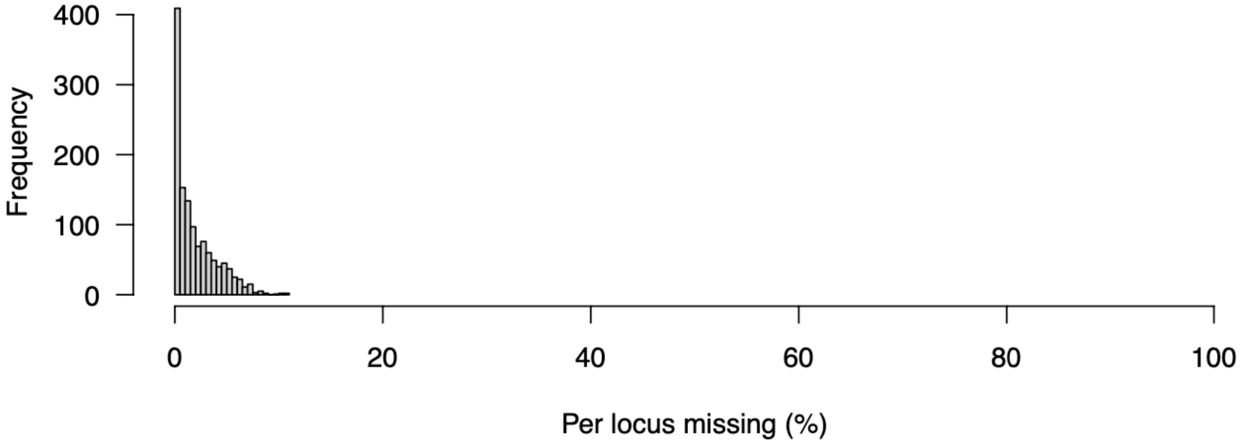


(B)

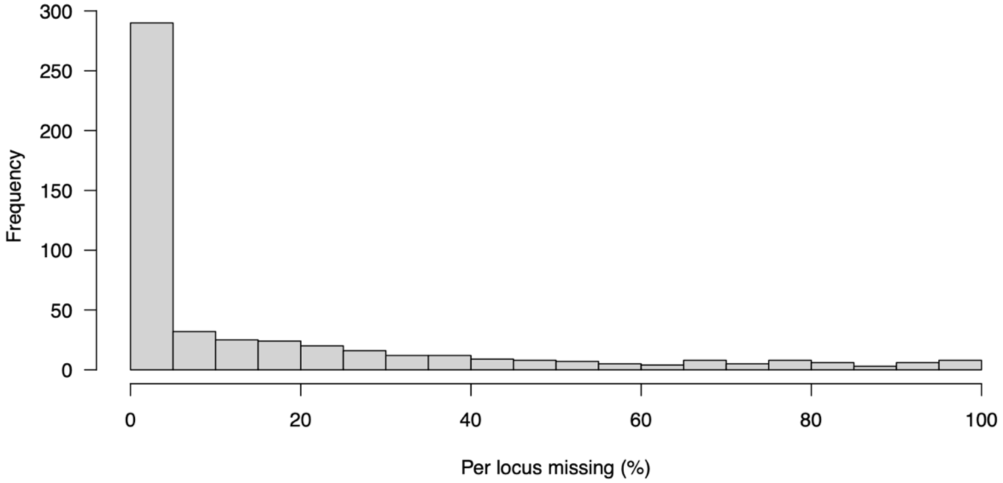


**Figure S2.** Percentage of per-locus missing data in the amplicon panel genotyped (A) using all *de novo* identified SNPs (n = 1,257 SNPs); and (B) hotspots only transferred to the chromosome-level assembly (n = 508 SNPs).

(A)

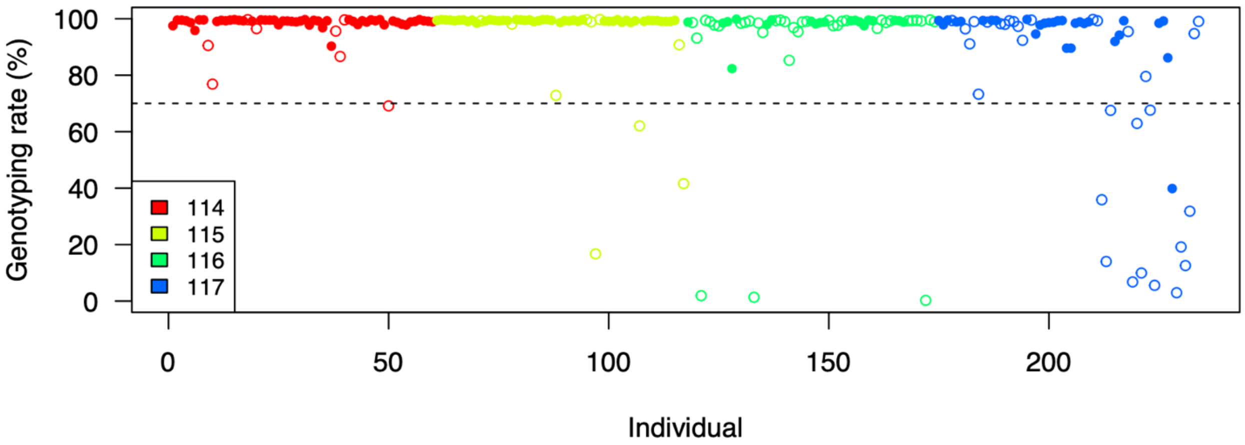


(B)

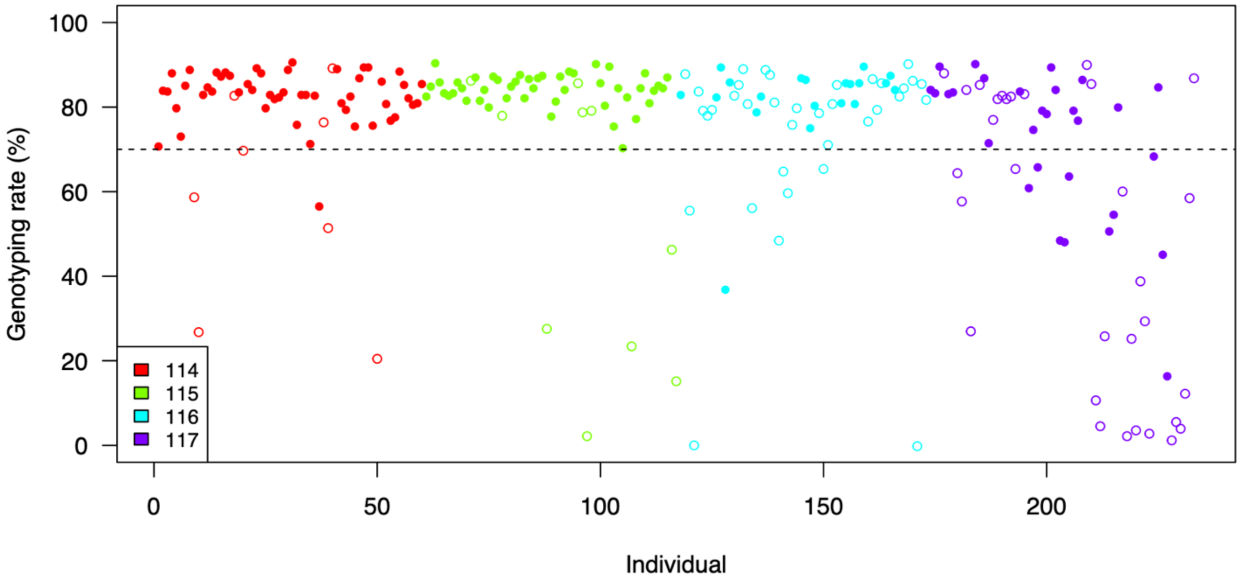


**Figure S3.** Genotyping rate (%) per sample using the amplicon panel for (A) *de novo* identified SNPs (n = 1,257 SNPs); and (B) hotspot-only SNPs transferred to the chromosome-level assembly (n = 508 SNPs). Families are indicated by colour, filled in circles indicate survivors whereas empty circles are mortalities.

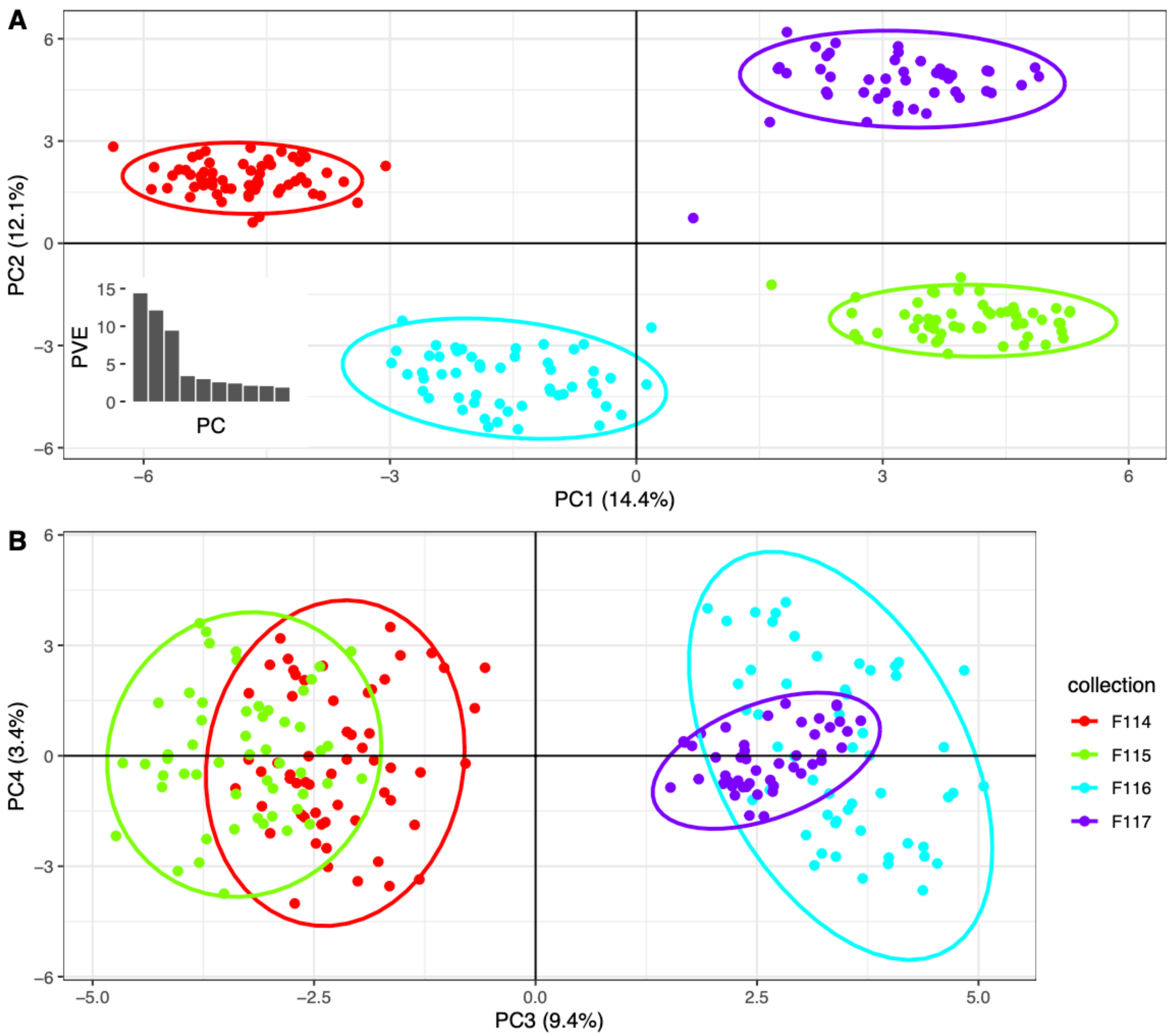


**Figure S4.** Principal component analysis (PCA) showing (A) PC1 and PC2 and (B) PC3 and PC4 clustering of samples by genotypes based on the amplicon panel *de novo* SNPs. Samples clustered by family, and the first three axes captured most of the variation.

**(A)**

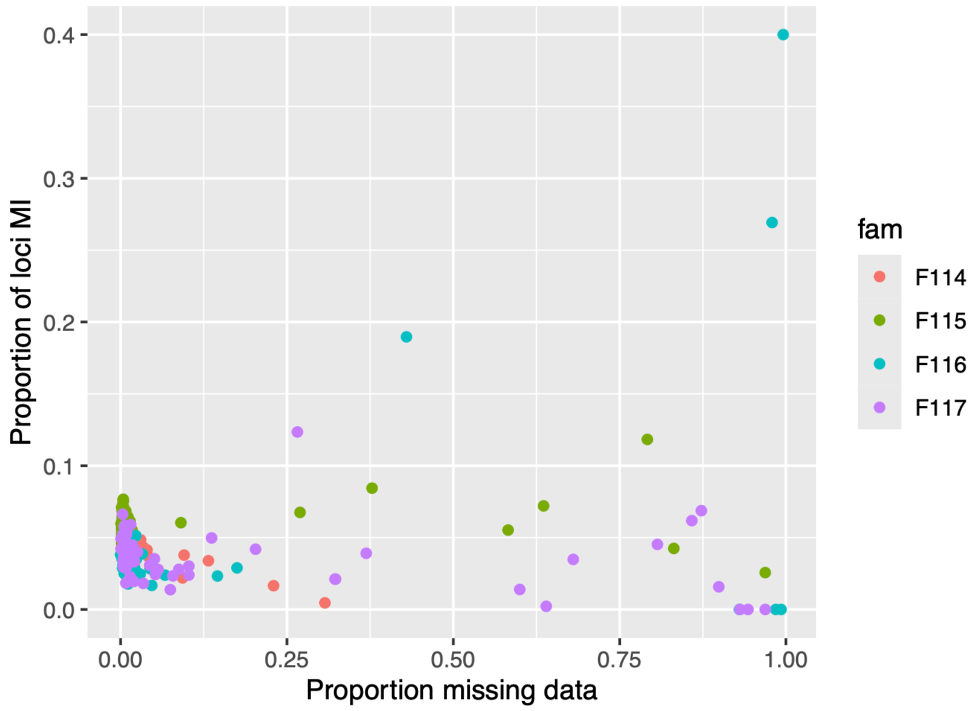


**(B)**

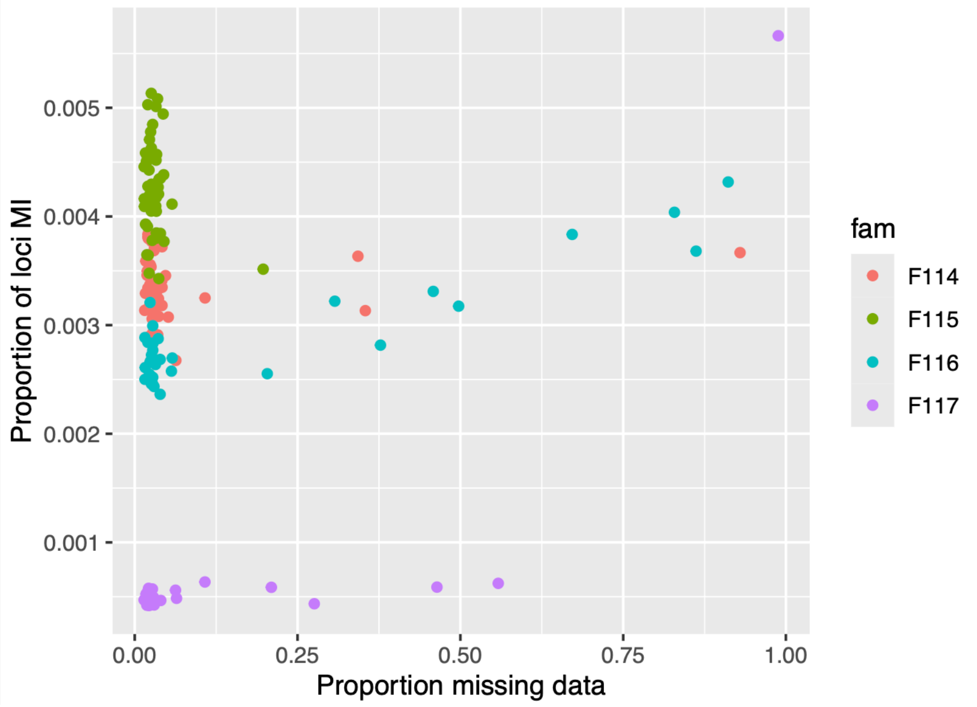


**Figure S5.** Per individual proportions of loci that exhibit Mendelian incompatibilities shown relative to the proportion of missing data per individual, and coloured by family in (A) the amplicon panel dataset (LD data; n = 240 individuals); and (B) the whole-genome resequencing dataset (HD data; n = 176 individuals).

### SUPPLEMENTAL TABLES

**Table S1.** Number and percentage of total variants retained through all filtering steps for whole-genome resequencing data (HD data) or amplicon panel *de novo* genotyping data (LD data). Both datasets were genotyped with all individuals together (HD data: n = 8 parents; 176 offspring; LD data: n = 8 parents; 240 offspring).

| Filtering stage | No. variants retained, HD | % retained, HD | No. variants retained, LD | % retained, LD |
|---|---|---|---|---|
| All variants | 58,941,933 | - | 218,925 | - |
| Retain variants > 5 bp from indel | 52,888,788 | 89.7% | 217,222 | 99.2% |
| Retain if missing in < 10% (HD) or 15% (LD) samples | 29,772,551 | 50.5% | 3,673 | 1.7% |
| SNP only | 27,816,124 | 47.2% | 2,810 | 1.3% |
| SNP quality ≥ 99 in at least one sample | 24,062,856 | 40.8% | 2,666 | 1.2% |
| Average depth across samples > 10 reads | 6,811,239 | 11.6% | 2,666 | 1.2% |
| Retain biallelic SNPs only | 6,245,592 | 10.6% | 2,624 | 1.2% |
| Per sample, per genotype set as missing if depth < 5 or > 100 (HD), or depth < 10 or > 100K (LD); or if genotype quality < 20.<br><br>Retain if missing in < 10% (HD) or 15% (LD) samples | 782,033 | 1.3% | 1,501 | 0.7% |
| MAF < 0.05 | **424,951** | 0.7% | **1,257** | 0.6% |

**Table S2.** Per family numbers of loci with complete genotypes (i.e., no missing data) in parents that were monomorphic or polymorphic in parents or offspring, and the number and percentage of complete genotypes that were unexpectedly polymorphic in offspring as the locus was monomorphic in the respective parents.

| | Parents | | | | Offspring | | | |
|---|---|---|---|---|---|---|---|---|
| | # loci, complete | # mono-morphic | # poly-morphic | % poly-morphic | # mono-morphic | # poly-morphic | % poly-morphic | Unexpected polymorphic |
| F114 | 402,019 | 210,115 | 191,904 | 47.7 | 204,824 | 197,195 | 49.1 | 5,302 (2.7%) |
| F115 | 411,022 | 206,271 | 204,751 | 49.8 | 199,808 | 211,214 | 51.4 | 6,472 (3.1%) |
| F116 | 399,274 | 230,313 | 168,961 | 42.3 | 226,111 | 173,163 | 43.3 | 4,253 (2.5%) |
| F117 | 231,194 | 122,593 | 108,601 | 47.0 | 122,093 | 109,101 | 47.2 | 500 (0.5%) |
| AVG. | 360,877 | 192,323 | 168,554 | 46.7 | 188,209 | 172,668 | 47.8 | 4132 (2.2%) |
