## Supplementary figures and images for "Pedigree-based genome-wide imputation using a low-density amplicon panel for the highly polymorphic Pacific oyster *Crassostrea* (*Magallana*) *gigas*"

### Additional File S2

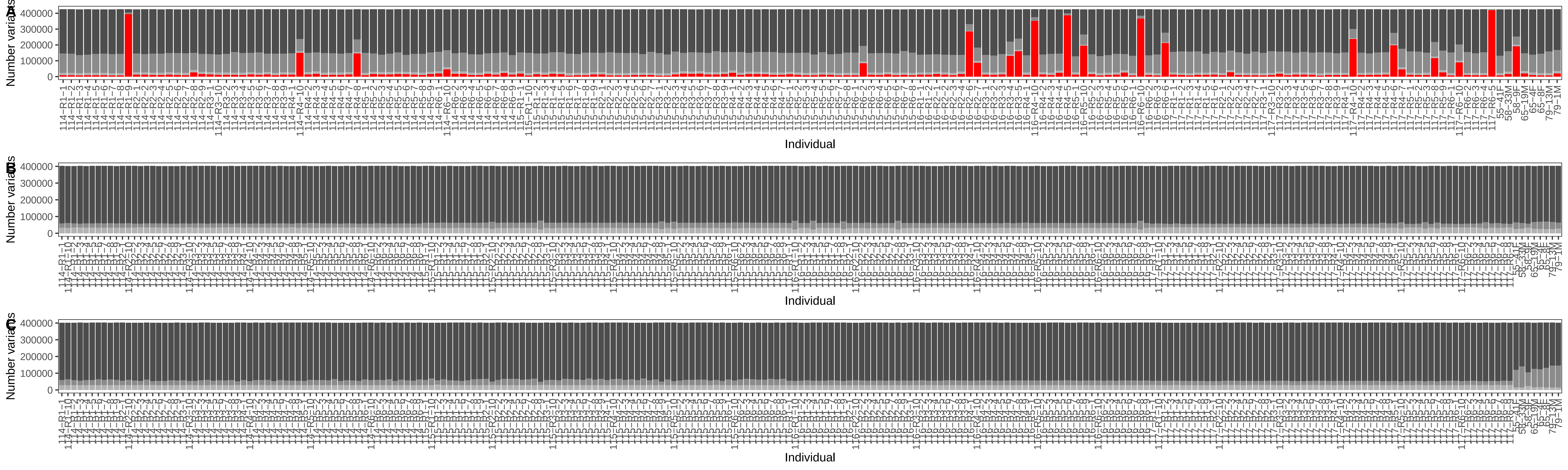
